# A Pif1-dependent threshold separates DNA double-strand breaks and telomeres

**DOI:** 10.1101/078162

**Authors:** Jonathan Strecker, Daniel Durocher

## Abstract

The natural ends of chromosomes resemble DNA double-strand breaks (DSBs) and telomeres are therefore necessary to prevent recognition by the DNA damage response. The enzyme telomerase can also generate new telomeres at DSBs, resulting in the loss of genetic information distal to the break. How cells deal with different DNA ends is therefore an important decision. One critical point of regulation is to limit telomerase activity at DSBs and this is primarily accomplished in budding yeast by the telomerase inhibitor Pif1. Here we use Pif1 as a sensor to gain insight into the cellular decision at DSB ends with increasing telomeric character. We uncover a striking transition point in which 34 bp of telomeric (TG_1-3_)_n_ repeat sequence is sufficient to render a DNA end insensitive to Pif1, thereby facilitating extension by telomerase. This phenomenon is unlikely to be due to Pif1 modification and we propose that Cdc13 confers a unique property to the TG_34_ end that prevents Pif1 action. We identify novel Cdc13 mutations that resensitize DNA ends to Pif1 and discover that many Cdc13 telomerase-null mutations are dependent on Pif1 status. Finally, the observed threshold of Pif1 activity recapitulates several properties of both DSBs and telomeres and we propose that this is the dividing line between these entities.

## Introduction

A fundamental question in chromosome biology is how cells differentiate between spontaneous DNA double-strand breaks (DSBs) and telomeres, the natural ends of chromosomes. The failure to properly deal with each end has severe consequences for the cell and the inappropriate repair of telomeres can lead to chromosome fusions and mitotic breakage. Similarly, the activity of telomerase at DSBs can generate new telomeres, at a cost of the genetic information distal to the break. Telomere addition has been observed in a variety of species (Biessmann et al., 1990; Fouladi et al., 2000; Kramer and Haber, 1993) and has been linked to human disorders involving terminal deletions of chromosome 16 (Wilkie et al., 1990) and 22 (Wong et al., 1997). While DSBs and telomeres reflect extreme positions on the spectrum, a continuum of DNA ends exist between them including critically short telomeres and DSBs occurring in telomeric-like sequence. All of these require a decision: should the end be repaired or elongated by telomerase?

The budding yeast *Saccharomyces cerevisiae* has been a key model to study mechanisms of genomic stability and whose telomeres consist of 300±75 bp of heterogeneous (TG_1-3_)_n_ repeats (Zakian, 1996). One way to deal with the DNA end problem would be to exclude DSB repair proteins from telomeres, but paradoxically this is not the case as repair complexes including MRX and Ku have important roles at telomeres (Lydall, 2009). The activity of telomerase must therefore be tightly regulated at DSB sites and this is accomplished in budding yeast by the telomerase inhibitor Pif1 (Schulz and Zakian, 1994; Zhou et al., 2000). Pif1 has both mitochondrial and nuclear isoforms encoded from separate translational start sites; mutation of the second start site in the *pif1-m2* mutant abolishes the nuclear isoform (Schulz and Zakian, 1994) resulting in longer telomeres and a 240-fold increase in telomere addition at DSBs (Bochman et al., 2010; Myung et al., 2001). Pif1 is a helicase that preferentially unwinds RNA-DNA hybrids in vitro (Boulé et al., 2005) and is thought to remove the TLC1 telomerase RNA template from telomeres (Li et al., 2014; Phillips et al., 2015); however, it is unclear whether Pif1 performs the same function at DSBs. Arguing against this idea is the observation that telomere addition events do not preferentially occur at TLC1 binding sites in the absence of Pif1 (Putnam et al., 2004).

Interestingly, Pif1 is able to distinguish between DSBs and telomeres as a *pif1-4A* mutant affects telomere addition frequency but not telomere length (Makovets and Blackburn, 2009), making Pif1 an attractive candidate to control the fate of DNA ends. Previous work in our lab revealed that Pif1 suppresses telomere addition at DNA ends containing 18 bp of (TG_1-3_)_n_ telomeric repeats (referred to as TG_18_), but has no effect at the TG_82_ end (Zhang and Durocher, 2010). This result suggests that the TG_82_ substrate is interpreted by the cell to be a short telomere and is allowed to elongate in a manner uninhibited by Pif1. We sought to investigate the molecular basis of this DNA end-fate decision using the activity of Pif1 as a cellular sensor.

## Results

### Identification of a Pif1 threshold at DNA ends

To characterize the dividing line between DSBs and telomeres we used a genetic system in which galactose-inducible HO endonuclease can be expressed to create a single DSB at the *ADH4* locus on Chr VII-L (Diede and Gottschling, 1999; Gottschling et al., 1990). By placing different lengths of telomeric (TG_1-3_)_n_ sequence immediately adjacent to the HO cut site one can study the fate of DNA ends using two readouts: a genetic assay for telomere addition based on the loss of the distal *LYS2* marker, and by Southern blotting to monitor the length of the DNA end (**Figure 1ab**). The HO cut site in this system contributes one thymine nucleotide to the inserted telomeric seed, accounting for a one base pair discrepancy from prior reports. As previous work indicated that Pif1 is active at TG_18_, but not TG_82_ (Zhang and Durocher, 2010), we first constructed strains containing 34, 45, 56, and 67 bp of telomeric repeats in both wild-type and *pif1-m2* cells (see **Supplementary Table 1** for all TG repeat sequences). We observed similar rates of telomere addition at all DNA ends in both backgrounds, indicating that 34 bp of telomeric repeat is sufficient to render a DNA end insensitive to Pif1 (**Figure 1c**). To account for variations in HO cutting efficiency and the propensity to recruit telomerase at each DNA end, we also normalized telomere addition frequency to *pif1-m2* cells to provide a clear readout of Pif1 activity (**Figure S1a**). Analysis of DNA ends by Southern blot also revealed robust telomere addition at the TG_34_ substrate in *PIF1* cells mirroring the results of the genetic assay (**Figure 1d**).

**Figure 1.**
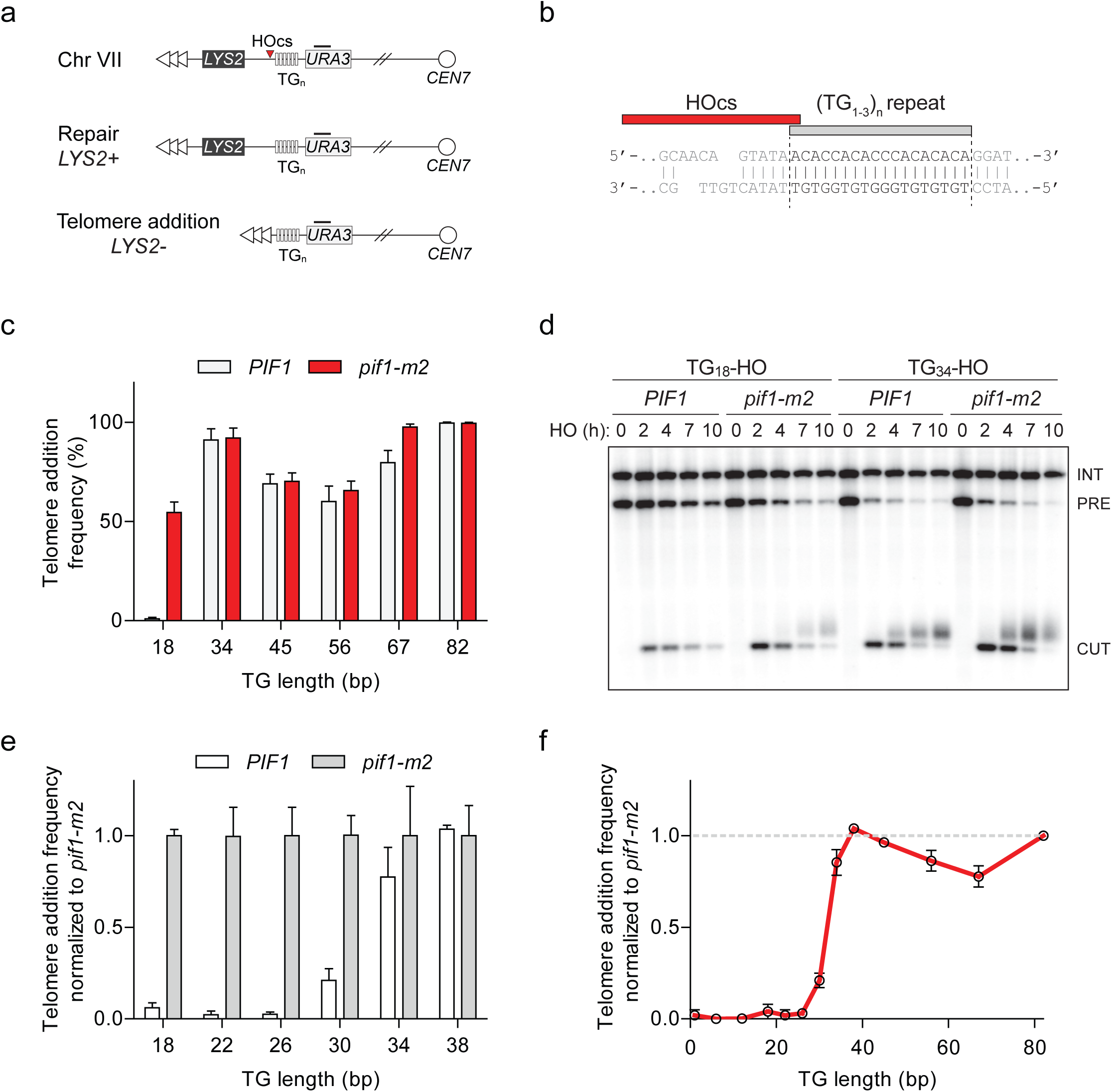
Characterization of Pif1 activity at DNA ends reveals a DSB-telomere transition. **a**, Schematic of a system to study the fate of DNA ends. Telomeric repeats are placed adjacent to an HO cut site (HOcs) at the *ADH4* locus on Chr VII. Telomere addition can be measured using a genetic assay based on the loss of the distal *LYS2* gene as measured by resistance to α-aminoadipic acid. Southern blotting with a probe complementary to *URA3* (black bar) allows for visualization of DNA end stability. **b**, Sequence of the TG_18_ substrate and the overhang produced by the HO endonuclease. The C-rich strand runs 5’ to 3’ towards the centromere and is resected following DSB induction to expose a 3’ G-rich overhang. **c**, Telomere addition frequency at DNA ends containing 18-82 bp of TG sequence. Data represents the mean ± s.d. from a minimum of n=3 independent experiments. See Supplementary Table 1 for the sequences of all DNA ends. **d**, Southern blot of DNA ends containing TG_18_ and TG_34_ ends in wild-type and *pif1-m2* cells following HO induction. A *URA3* probe was used to label the *ura3-52* internal control (INT) and the *URA3* gene adjacent to the TG_n_-HO insert (PRE) which is cleaved by HO endonuclease (CUT). Newly added telomeres are visualized as a heterogeneous smear above the CUT band. **e**, Telomere addition frequency normalized to *pif1-m2* cells at DNA ends containing 18-38 bp of TG sequence. Data represents the mean ± s.d. from n=3 independent experiments. **f**, Summary of telomere addition frequency normalized to *pif1-m2* across the spectrum of TG repeat substrates.

The standard genetic telomere addition assay includes a nocodazole arrest before DSB induction as telomerase is active in S and G2 phase (Diede and Gottschling, 1999). However, asynchronously dividing cells also exhibited a similar phenotype at the TG_18_ and TG_34_ ends (**Figure S1b**). To exclusively study telomere addition by telomerase and not through genetic recombination, telomere addition strains also harbour a *rad52Δ* mutation. The addition of *RAD52* in this assay reduced telomere addition at the TG_18_ in *PIF1* cells but had no impact on the behavior of Pif1 at the TG_34_ substrate (**Figure S1b**).

To further refine the threshold, we added 4 bp increments of TG repeat sequence to the centromeric side of the TG_18_ substrate yielding strains with 22, 26, 30, 34, and 38 bp of telomeric repeats. Importantly, with the exception of length, these strains contain the same DNA sequence and share a common distal end. Analysis of telomere addition revealed that Pif1 is active at DNA ends up to TG_26_ while the frequency of telomere addition increased at the TG_30_ end and beyond (**Figure 1e**). As telomeric repeats are heterogeneous in nature, we next determined if this phenotype is dependent on the particular DNA sequence. We selected three sequences at random from *S. cerevisiae* telomeric DNA and constructed strains with DNA ends containing either 26 or 36 bp of each sequence. Consistent with our initial observations, telomere addition was inhibited by Pif1 at all TG_26_ ends, while the corresponding TG_36_ ends resulted in telomere addition in the presence of Pif1 (**Figure S1cd**).

Visualization of the combined genetic assay results across different lengths of TG repeat substrates revealed a striking transition with regards to Pif1 function (**Figure 1f**). By using Pif1 as a cellular sensor we propose that the 26 to 34 bp window of telomeric sequence is the dividing line between what the cell interprets to be a DSB, and what is considered to be a critically short telomere. These data suggest that DNA ends containing 34 bp or more of telomeric DNA are allowed to elongate in a manner unimpeded by Pif1 and we herein refer to this phenomenon as the DSB-telomere transition.

### Pif1 is not inhibited by DNA damage kinases

One attractive mechanism for the observed DSB-telomere transition is that Pif1 might be inactivated at DNA ends containing longer telomeric repeats. Prime candidates for this regulation include the central DNA damage kinases including Mec1, Tel1, and Rad53. Previous work has identified that Tel1 promotes telomerase-mediated extension of the TG_82_ end (Frank et al., 2006), and targets short telomeres for elongation (Sabourin et al., 2007). As these results raised the possibility that Tel1 antagonizes Pif1, we deleted *TEL1* in both wild-type and *pif1-m2* backgrounds and followed the fate of the TG_82_ DNA end by Southern blotting. Although telomere addition was reduced in *tel1Δ* cells, we observed a similar reduction in *tel1Δ pif1-m2* cells indicating that *TEL1*and *PIF1* function in separate pathways (**Figure 2ab**). Consistent with this observation, the loss of *TEL1* did not affect the DSB-telomere transition at the TG_18_ and TG_34_ DNA ends (**Figure 2c**). Loss of *MEC1* and *RAD53* also failed to inhibit telomerase in a Pif1-specific manner at the TG_82_ end (**Figure S2a-d**). Pif1 contains five consensus S/T-Q Mec1 and Tel1 phosphorylation sites in Pif1; however, their mutation in the *pif1-5AQ* allele (S148A/S180A/T206A/S707A/T811A) also did not decrease telomere addition at the TG_34_ end (**Figure 2d**).

**Figure 2.**
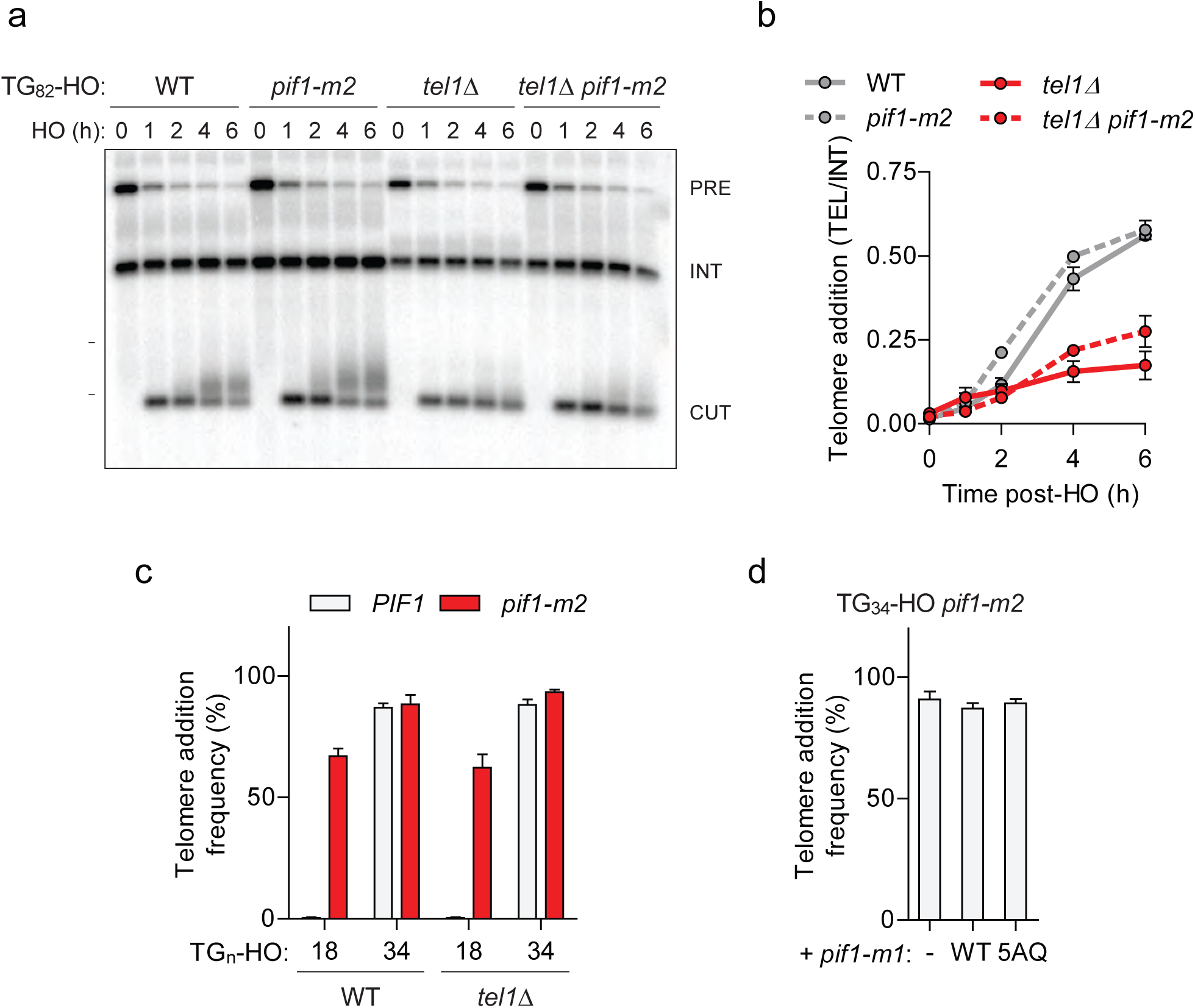
Pif1 is not inactivated by Tel1 at short telomeres. **a, b**, Southern blot of the TG_82_ DNA end following HO-induction in wild-type (WT) and *pif1-m2* cells without or with a *TEL1* deletion. An *ADE2* probe was used to label the *ade2Δ1* internal control (INT) and the *ADE2* gene adjacent to the TG_n_-HO insert (PRE) which is cleaved by HO endonuclease (CUT). Quantification of the newly added telomere signal (**b**) calculated by subtracting the background signal present prior to HO-induced and by normalizing to the INT control. Data represents the mean ± s.d. from n=2 independent experiments. **c**, Telomere addition frequency at the TG_18_ and TG_34_ DNA ends in *tel1Δ* mutants. Data represents the mean ± s.d. from n=3 independent experiments. **d**, Telomere addition frequency at the TG_34_ DNA end in *pif1-m2* cells (-) and cells expressing a wild-type (WT) or *pif1-5AQ* (S148A, S180A, T206A, S707A, T811A) nuclear-specific *pif1-m1* allele. Data represents the mean ± s.d. from n=3 independent experiments.

As Pif1 might be regulated though unanticipated post-translational modifications or protein interactions we performed a *PIF1* PCR mutagenesis screen to identify gain-of-function mutations that inhibit telomere addition at the TG_82_ end but we were unable to recover any mutants (**Figure S2ef**). Together these data challenge the hypothesis that Pif1 is inactivated at the TG_34_ and TG_82_ DNA ends, so we next considered alternative explanations for the observed DSB-telomere transition.

### Artificial telomerase recruitment does not outcompete Pif1

A simple explanation for the DSB-telomere transition is that longer telomeric repeats might have an increased ability to recruit telomerase. If correct, this model predicts that artificially increasing telomerase recruitment to the TG_18_ end might be sufficient to overcome Pif1 inhibition. Since the primary mechanism of telomerase recruitment involves an interaction between the DNA binding protein Cdc13 and the Est1 telomerase subunit (Nugent et al., 1996; Pennock et al., 2001), we expressed Cdc13-Est1 and Cdc13-Est2 fusion proteins (Evans and Lundblad, 1999) to test this possibility. In agreement with previous work, expression of both fusions resulted in greatly elongated telomeres (**Figure 3a**); however, they did not increase telomere addition at the TG_18_ DNA end in the presence of Pif1 (**Figure 3b**). To test whether the Cdc13-Est1 fusion protein is able to bind and extend the TG_18_ substrate, we repeated the genetic assays in *est1Δ* cells expressing a Cdc13-Est1 fusion containing the *est1-60* mutation (K444E) which disrupts the interaction of Est1 with endogenous Cdc13 (Pennock et al., 2001). Telomerase extension in these *est1Δ* cells must therefore arise from the ectopic construct we observed that Cdc13-Est1^K444E^ can extend the TG_18_ end only in the absence of *PIF1* (**Figure 3b**). Together these data indicate that Pif1 is able to effectively suppress telomere addition even in the presence of enhanced telomerase recruitment suggesting that increased telomerase recruitment to the TG_34_ end is unlikely to underpin the observed DSB-telomere transition.

**Figure 3.**
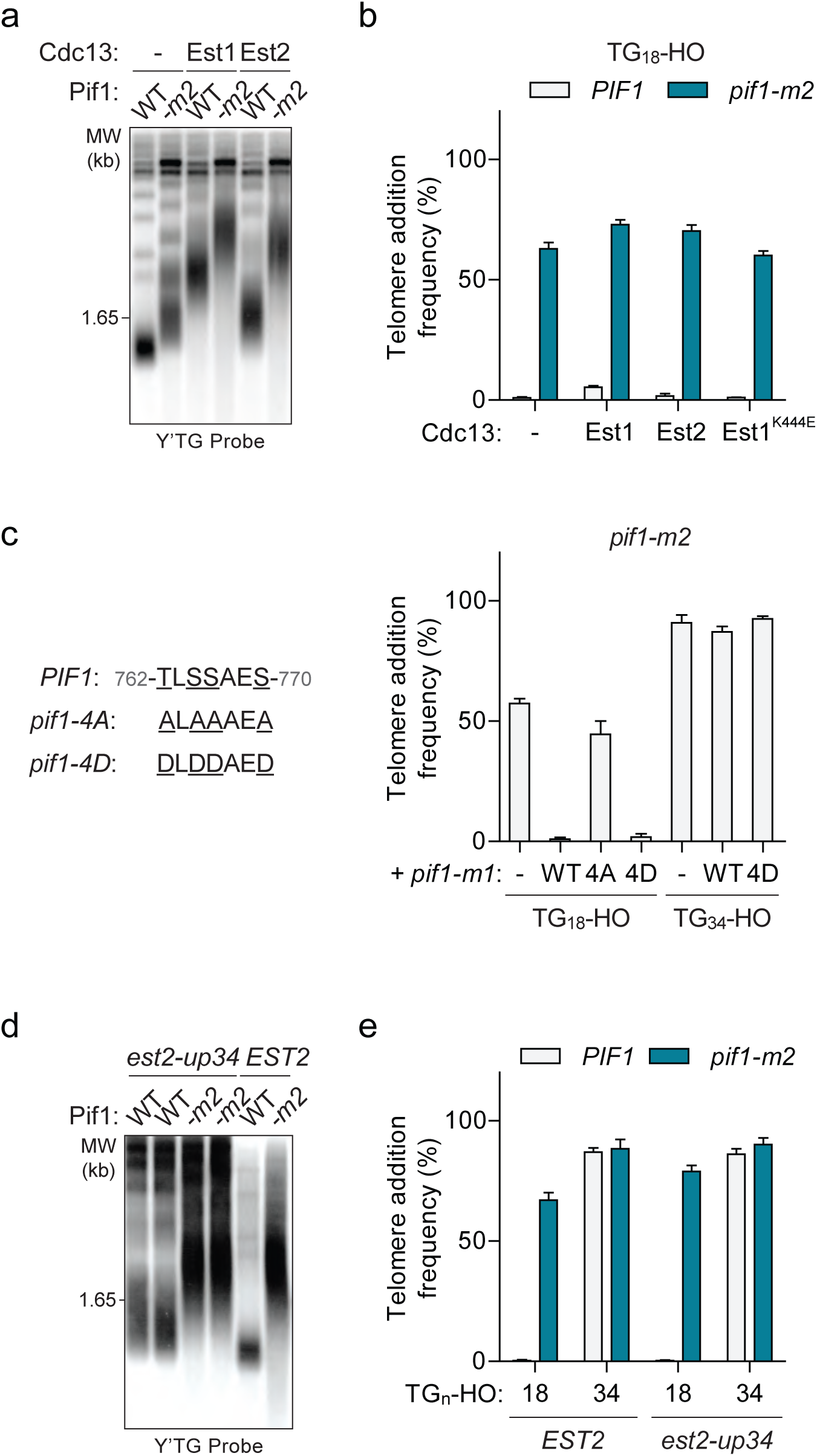
The DSB-telomere transition recapitulates the differential regulation of Pif1. **a**, Southern blot for telomere length in TG_18_-HO wild-type (WT) and *pif1-m2* cells harbouring an empty plasmid (-) or expressing plasmid-based Cdc13-Est1 or Cdc13-Est2 fusions. Cells were passaged for approximately 75 generations before genomic DNA extraction. A Y’TG probe was used to label telomeric sequences. **b**, Telomere addition frequency of the cells described in panel **a**, and *est1Δ* strains expressing a *cdc13-est1-60* (K444E) fusion. Data represents the mean ± s.d. from n=3 independent experiments. **c**, Telomere addition frequency at the TG_18_ and TG_34_ DNA ends in *pif1-m2* cells (-) and cells expressing a wild-type (WT), *pif1-4A* (T763A/S765A/S766A/S769A), or *pif1-4D* (T763A/S765A/S766A/S769A) nuclear specific *pif1-m1* allele. Data represents the mean ± s.d. from n=3 independent experiments. **d**, Southern blot for telomere length in *PIF1* (WT) and *pif1-m2* cells combined with or without the *est2-up34* mutation. Cells were passaged for approximately 75 generations before genomic DNA extraction and a Y’TG probe was used to label telomere sequences. **e**, Telomere addition frequency at the TG_18_ and TG_34_ DNA ends in *PIF1* and *pif1-m2* cells with or without the *est2up-34* mutantion. Data represents the mean ± s.d. from n=3 independent experiments.

### The DSB-telomere transition recapitulates the differential regulation of Pif1

While we previously hypothesized that Pif1 might be inhibited at DNA ends that resemble telomeres, an alternative possibility is that Pif1 is only activated at DNA ends with short tracts of telomeric sequence. Consistent with this model, Pif1 is reported to be phosphorylated after DNA damage in a Mec1-Rad53-Dun1-dependent manner and further characterization of this activity led to the identification of the *pif1-4A* mutant (T763A/S765A/S766A/S769A) that is unable to inhibit telomere addition at DSBs (Makovets and Blackburn, 2009). Importantly, mimicking phosphorylation with the *pif1-4D* allele can restore Pif1 activity (Makovets and Blackburn, 2009). We first confirmed the function of these mutants at the TG_18_ DNA end by integrating variants of the nuclear specific *pif1-m1* allele at the *AUR1* locus in *pif1-m2* cells (**Figure 3c**). If Pif1 phosphorylation only occurs at DNA ends with short lengths of telomeric repeats, such as TG_18_, then mimicking phosphorylation may be sufficient to inhibit telomere addition at DNA ends with longer telomeric repeats. Contrary to this prediction, the *pif1-4D* mutant did not restore Pif1 activity at the TG_34_ DNA end (**Figure 3c**) indicating that phosphorylation of these sites does not regulate the DSB-telomere transition.

Several lines of evidence indicate that Pif1 functions differently at DSBs and telomeres. First, the *pif1-4A* mutation affects telomere addition at DSBs, but not telomere length (Makovets and Blackburn, 2009). The inability of the *pif1-4D* allele to inhibit telomerase at TG_34_ therefore provides indirect evidence that this DNA end is interpreted by the cell as a short telomere. A second mutation that affects Pif1 activity has also been identified: the *est2-up34* mutation, which affects the finger domain of the telomerase reverse transcriptase subunit (Eugster et al., 2006). Interestingly, the *est2-up34* mutant results in over-elongated telomeres in wild-type but not *pif1-m2* cells, indicating that the *est2-up34* allele can at least partially bypass Pif1 inhibition (Eugster et al., 2006). To test if this holds true at DSBs we generated the *est2-up34* mutation in strains with a TG_18_DNA end. Although we observed increased telomere length in *PIF1 est2-up34* cells (**Figure 3d**), telomere addition was not increased (**Figure 3e**), indicating that the *est2-up34* mutation can bypass Pif1 function at telomeres but not at DSBs. Together these data support the idea that Pif1 possesses distinct functions at DSBs and telomeres, and that these differences are recapitulated in the TG_18_ and TG_34_ DNA ends on either side of the DSB-telomere transition.

### Investigating the molecular trigger of the DSB-telomere transition

Since our attempts thus far failed to identify a modification of Pif1 that would explain the DSB-telomere transition, we next focused on whether a property of the DNA end facilitates or blocks Pif1 activity. Attractive candidates included the MRX and Ku complexes which are rapidly recruited to DNA ends and function in both DSB repair and telomere maintenance. The loss of either complex, however, did not effect either side of the DSB-telomere transition (**Figure 4a**).

**Figure 4.**
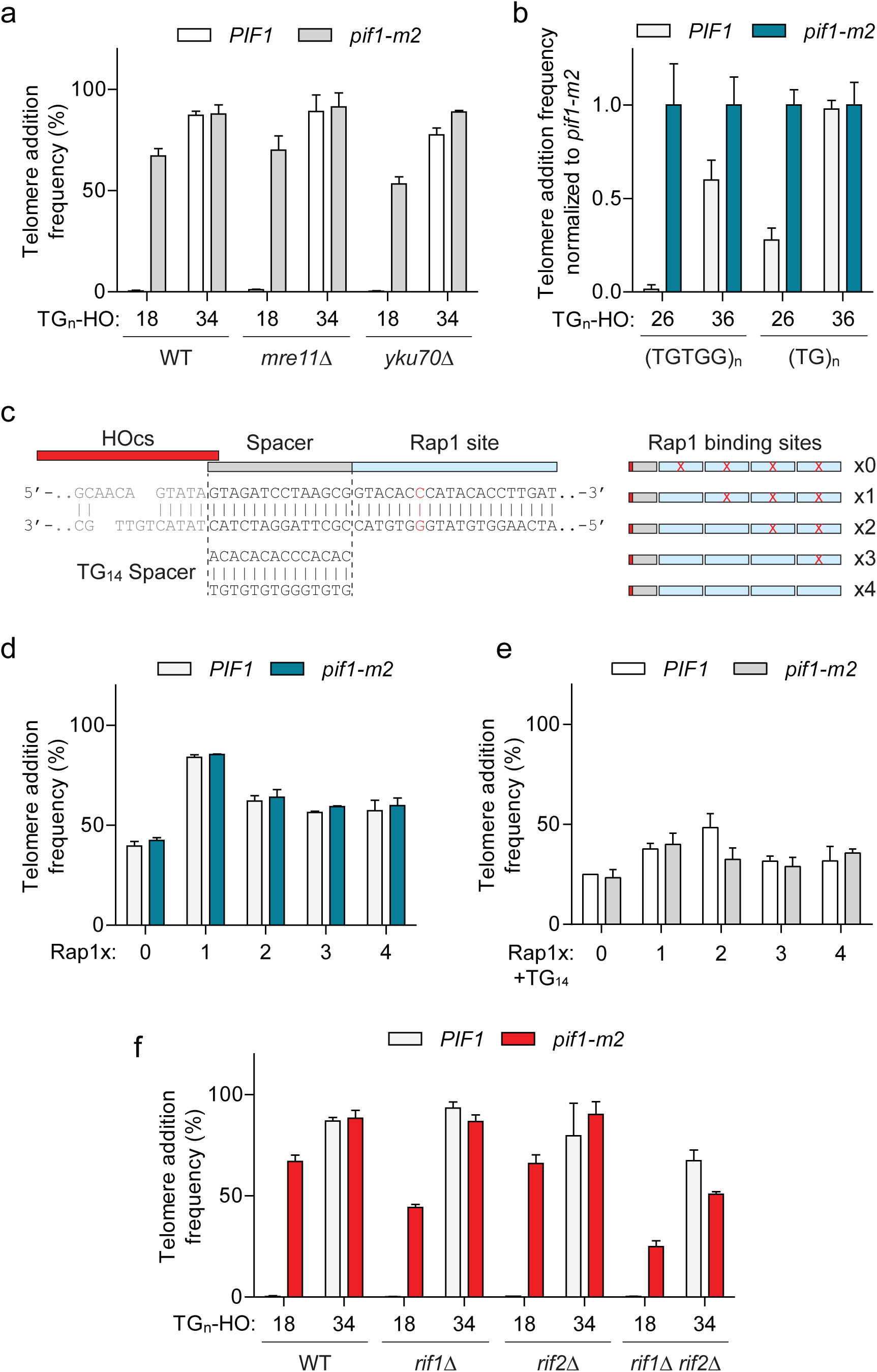
The DSB-telomere transition does not require Rap1. **a**, Telomere addition frequency at the TG_18_ and TG_34_ DNA ends in *mre11Δ* and *yku70Δ* mutants. Data represents the mean ± s.d. from n=3 independent experiments. **b**, Telomere addition frequency normalized to *pif1-m2* cells at DNA ends containing 26 bp or 36 bp of either (TGTGG)_n_ or (TG)_n_ repeats. Data represents the mean ± s.d. from n=3 independent experiments. **c**, Schematic of an array of Rap1 binding sites adjacent to the HO cut site on Chr VII with either a non-telomeric 14 bp spacer (upper) or a TG_14_ spacer (lower). Rap1 binding is disrupted by a cytosine to guanine mutation in the C-rich strand (position highlighted in red) to yield arrays of zero to four functional Rap1 sites. **d, e**, Telomere addition frequency at the Rap1 arrays described in panel **c** with a non-telomeric spacer (**d**) or TG_14_ spacer (**e**) between the HO cut site and the first Rap1 binding site. Data represents the mean ± s.d. from n=3 independent experiments. **f**, Telomere addition frequency at the TG_18_ and TG_34_ DNA ends in *rif1Δ, rif2Δ*, and *rif1Δ rif2Δ* double mutants. Data represents the mean ± s.d. from n=3 independent experiments.

Budding yeast telomeres are bound by two specialized proteins, Rap1 and Cdc13, and the binary nature of the DSB-telomere transition suggests that the discrete binding of either protein may trigger insensitivity to Pif1. As Cdc13 binds single-stranded DNA at the distal end of the telomere (Lin and Zakian, 1996; Nugent et al., 1996), an attractive prediction is that Rap1 bound to double-stranded telomeric DNA of longer TG repeats might inhibit Pif1. This model nicely correlates with the observed length of the DSB-telomere transition, as Cdc13 and Rap1 bind DNA sequences of 11 bp (Hughes et al., 2000) and 18 bp respectively (Gilson et al., 1993; Ray and Runge, 1999). Rap1 has also been previously shown to stimulate telomere addition (Grossi et al., 2001; Lustig et al., 1990; Ray and Runge, 1998).

Rap1 is an essential protein that binds the consensus DNA sequence of 5’-ACA<u>CCC</u>ATACACC-3’ containing an invariable CCC core (Graham and Chambers, 1994; Grossi et al., 2001; Wahlin and Cohn, 2000). Importantly, substitution of the middle cytosine to guanine in this motif abolishes Rap1 binding (Graham and Chambers, 1994; Grossi et al., 2001). To test whether Rap1 is required to bypass Pif1 activity at DNA ends we first generated synthetic telomeric sequences with strict (TGTGG)_n_ or (TG)_n_ repeats in both 26 bp and 36 bp lengths. Unlike natural telomeres, both sequences lack a CCC motif on the opposing strand. Despite these alternations, we still observed increased telomere addition at TG_36_ ends in wild-type cells (**Figure 4b**), suggesting that Rap1 binding is not required for this phenomenon.

In a second approach, we constructed an array of four consecutive Rap1 binding sites with a spacer adjacent to the HO cut site containing either 14 bp of non-telomeric DNA or a TG_14_ sequence (**Figure 4c**). Disrupting individual Rap1 binding sites with a cytosine to guanine substitution yielded arrays with zero to four functional binding sites (**Figure 4c**). Consistent with previous work, telomere addition was stimulated by a functional Rap1 binding site, but surprisingly all the tested arrays were insensitive to Pif1, including the array with no functional Rap1 binding sites (**Figure 4d**). Arrays with an adjacent TG_14_ spacer sequence displayed similar results with overall decreased rates of telomere addition possibly due to decreased HO cutting efficiency (**Figure 4e**). These results suggest that Rap1 binding is not required to bypass Pif1 activity and that additional features of the arrays are instead responsible. Finally, as telomere length regulation by Rap1 is coordinated through two downstream negative regulators of telomerase, Rif1 and Rif2 (Levy and Blackburn, 2004; Wotton and Shore, 1997), we asked whether these proteins are important for the DSB-telomere transition. Consistent with a Rap1-independent mechanism, telomere addition at the TG_34_ end was unaltered in *rif1Δ rif2Δ* mutants (**Figure 4f**).

### Cdc13 function influences the fate of DNA ends

Cdc13 binds a minimal 11 bp TG sequence through its canonical OB-fold DNA binding domain (Hughes et al., 2000) (**Figure 5a**) suggesting that three bound molecules of Cdc13 might render the TG_34_ DNA end insensitive to Pif1. The distinct switch of the DSB-telomere transition argues that this process might involve the assembly of a higher order protein complex. Alternatively, the Cdc13 N-terminal OB-fold domain (OB1) forms dimers (Mitchell et al., 2010; Sun et al., 2011) and can also bind telomeric ssDNA repeats in vitro of 37 and 43 bp, but not 18 and 27 bp (Mitchell et al., 2010), neatly matching our observed threshold. We hypothesized that Cdc13 dimerization and its unique N-terminal binding mode might allow longer DNA ends to bypass Pif1 and sought to test this idea by disrupting dimerization with the *cdc13-L91A* mutation (Mitchell et al., 2010). Consistent with this prediction, telomere addition at the TG_34_ end was inhibited by Pif1 in *cdc13-L91A* cells (**Figure 5b**); however, further investigation revealed a growth defect in these mutants that was suppressed by *pif1-m2* (**Figure 5c**). This result was reminiscent of the defective *cdc13-1* allele, which is also suppressed by loss of *PIF1* (Addinall et al., 2008; Downey et al., 2006). High copy plasmid expression of *cdc13-L91A* was able to rescue the growth defect, but also increased telomere addition at the TG_34_ substrate (**Figure 5b**) arguing that the initially observed defect in *cdc13-L91A* mutants was not solely due to impaired N-terminal dimerization.

**Figure 5.**
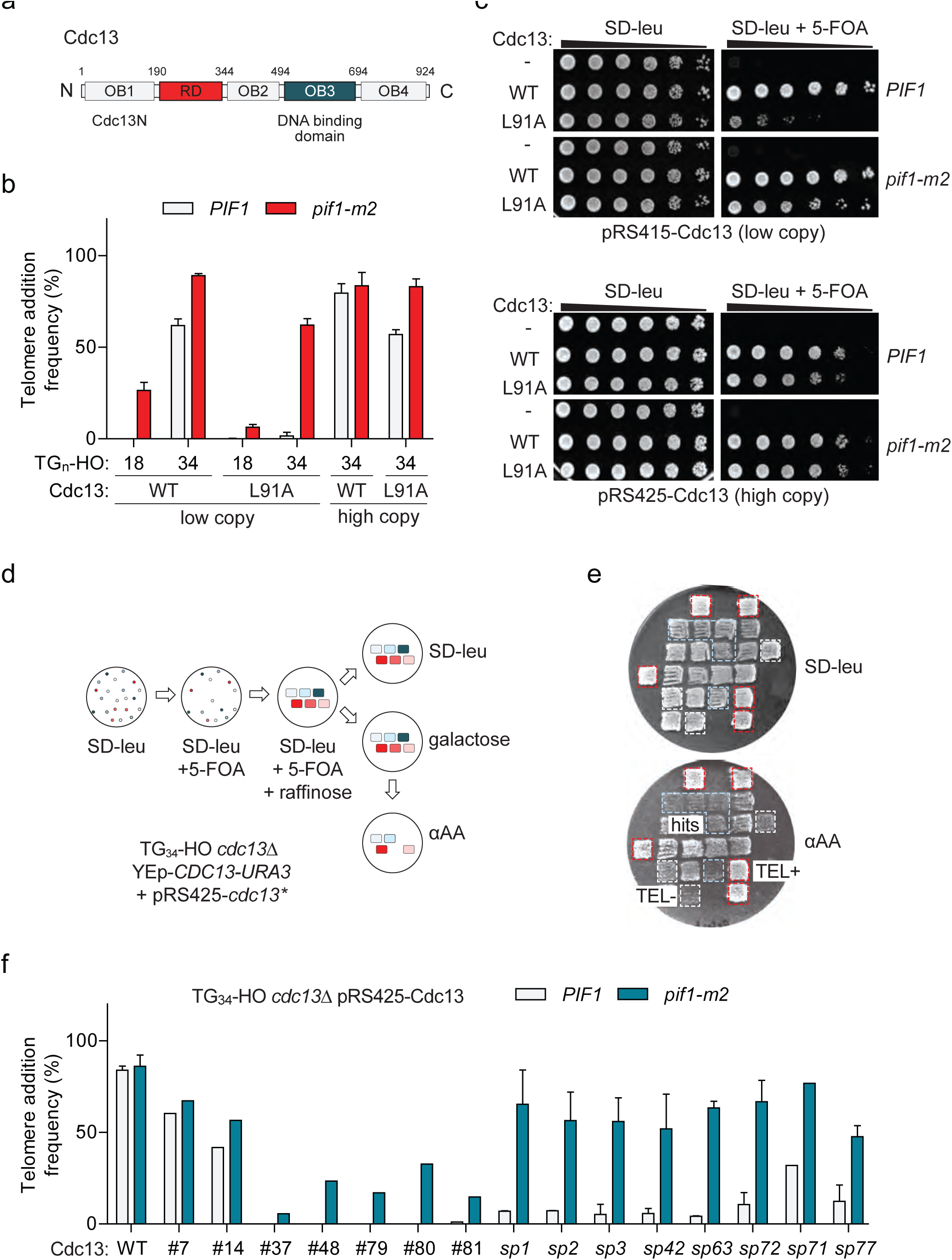
A genetic screen to identify Cdc13 mutants that prevent telomere addition at the TG_34_ end. **a**, Schematic of Cdc13 domain architecture consisting of four OB-fold domains (OB1-4) and a telomerase recruitment domain (RD). **b**, Telomere addition frequency at the TG_18_ and TG_34_ DNA ends in *cdc13Δ* cells expressing wild-type *CDC13* (WT) or *cdc13-L91A* from a low copy (pRS415) or high copy plasmid (pRS425). Data represents the mean ± s.d. from n=3 independent experiments. **c**, Spot assays to determine cell viability in *cdc13Δ* cells with a covering YEp-*CDC13-URA3* plasmid and pRS415- or pRS425-derived plasmids expressing wild-type Cdc13 (WT) or Cdc13-L91A. 5-fold serial dilutions of yeast cultures were grown on SD-leu as a control, and on SD-leu+5-FOA to determine viability in the absence of the covering plasmid. Plates were grown at 30°C for 2-3 days. **d**, Schematic of a screen in TG_34_ *cdc13Δ* cells using a plate-based genetic assay for telomere addition. Repaired mutant *cdc13* plasmids were selected on SD-leu and the covering YEp-*CDC13-URA3* removed by plating on 5-FOA before DSB induction. This step also eliminates all inviable *cdc13* mutations. Plates were incubated for 2-3 days at 30°C with the exception of galactose plates which were incubated for 4 hours. An agar plate was used to reduce cell number before final selection. **e**, Example re-testing plate from the screen. Cdc13 mutants that prevent telomere addition are identified by the inability to grow on media containing α-aminoadipate (αAA) (blue box), compared to positive control wild-type cells which add telomeres (red box) and TG_18_ cells that do not (white box). **f**, Telomere addition frequency at the TG_34_ DNA end in *PIF1* and *pif1-m2* cells in a *cdc13Δ* background expressing recovered pRS425-Cdc13 mutants from the screen. Data represents the mean ± s.d. from n=1 experiment for hits #7-81, and n=2 independent experiments for all *cdc13-sp* alleles.

We sought to more precisely dissect out a unique function of Cdc13 that blocks Pif1 and therefore performed a mutagenesis screen to identify *CDC13* alleles that have become sensitive to Pif1 activity (**Figure 5de**). Screening of approximately 6000 mutants led to the identification of fifteen hits that exhibited impaired telomere addition at the TG_34_ substrate. As this screen was performed in wild-type cells, we next determined if the mutations could support telomere addition in the absence of *PIF1*. Recovered plasmids were re-transformed into wild-type and *pif1-m2* cells, and analysis of telomere addition revealed two clones with minor phenotypes (#7 and 14), five clones with reduced telomere addition in both wild-type and *pif1-m2* cells (#37, 48, 79, 80, and 81), and eight clones in which telomere addition was impaired in wild-type cells but relatively unaffected in *pif1-m2* cells (#1, 2, 3, 42, 63, 71, 72, and 77) (**Figure 5f**). This observation suggested that the third group of Cdc13 mutations specifically sensitize the TG_34_ end to the activity of Pif1 and are herein referred to as *cdc13-sp* alleles (<u>s</u>ensitive to <u>P</u>if1).

DNA sequencing revealed an average of eleven amino acid substitutions per *cdc13-sp* allele and methodical mapping experiments led to the identification of causative amino acid substitutions in six of the eight *cdc13-sp* mutants (**Figure 6**, highlighted in red). Three alleles had contributions from multiple substitutions: I87N and Y758N in *cdc13-sp1*, H12R and F728I in *cdc13-sp72*, and E566V, N567, and Q583K in *cdc13-sp3* (**Figure 7a**). Cdc13-I87, like L91A, is also implicated in OB1 dimerization (Mitchell et al., 2010), again hinting that disrupting this function may restore Pif1 activity. The moderate telomere addition defect of Cdc13-I87N was likely only identified in the screen due to further exacerbation by the Y758N mutation (**Figure 7a**). The most important mutation in *cdc13-sp3* was identified to be Q583K with a minor contribution from E556V/N567D. Interestingly, all three residues are found in the canonical DNA binding domain, suggesting that weakening the association of Cdc13 with telomeric DNA can also sensitize the TG_34_ end to Pif1.

**Figure 6.**
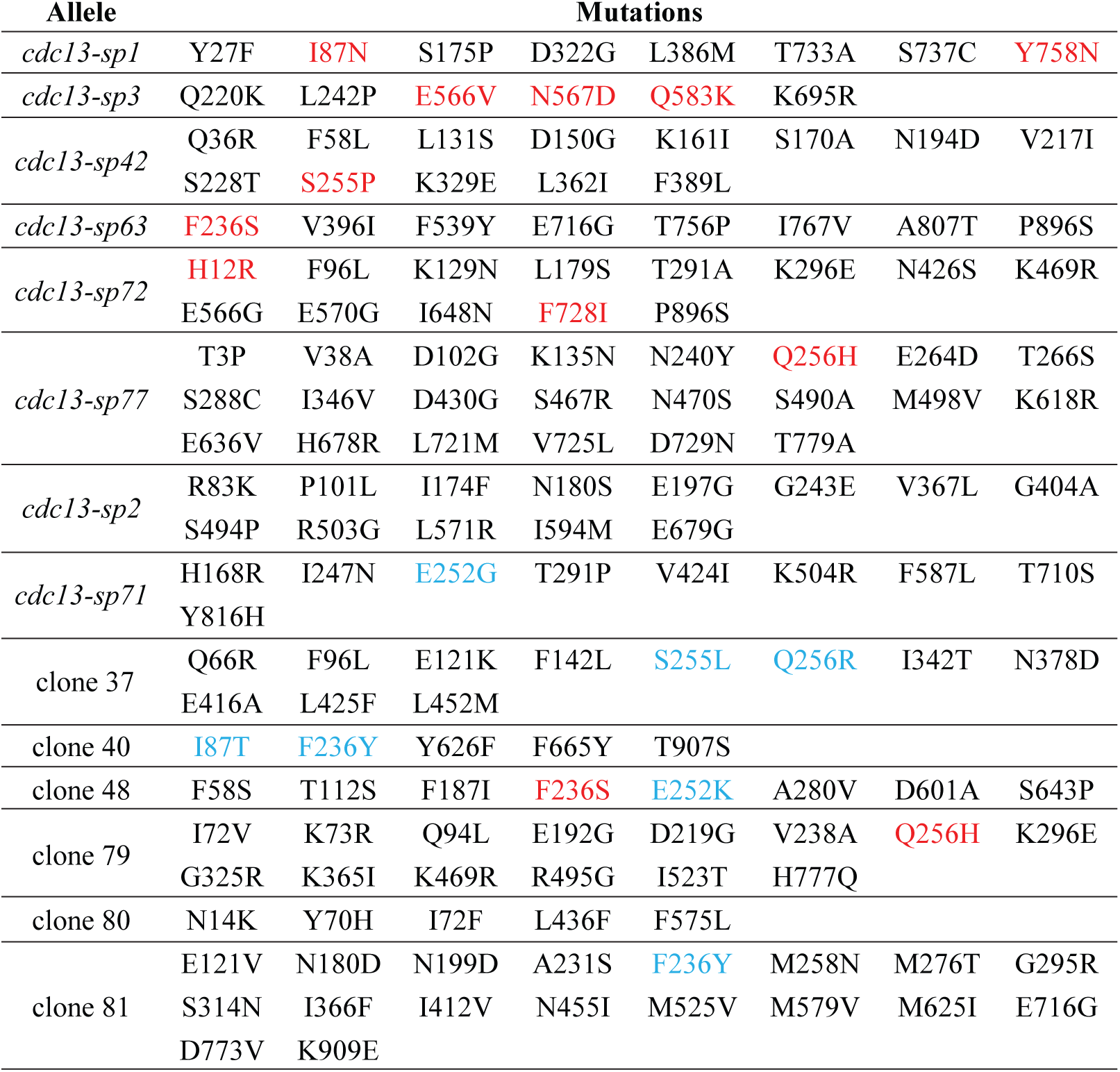
Mutations in *cdc13-sp* alleles. pRS425-*cdc13\** plasmids were recovered from cells grown on SD-leu media that were unable to grow on α-AA containing media. Cdc13 mutations were identified by plasmid sequencing. Mutations highlighted in red were identified by mapping experiments to determine which amino acid substitutions contribute to the mutant phenotype. Mutations highlighted in blue target important Cdc13 residues identified in this study or in previous work (Lendvay et al., 1996; Nugent et al., 1996) which are predicted to contribute to the defect although these exact substitutions were not specifically tested.

**Figure 7.**
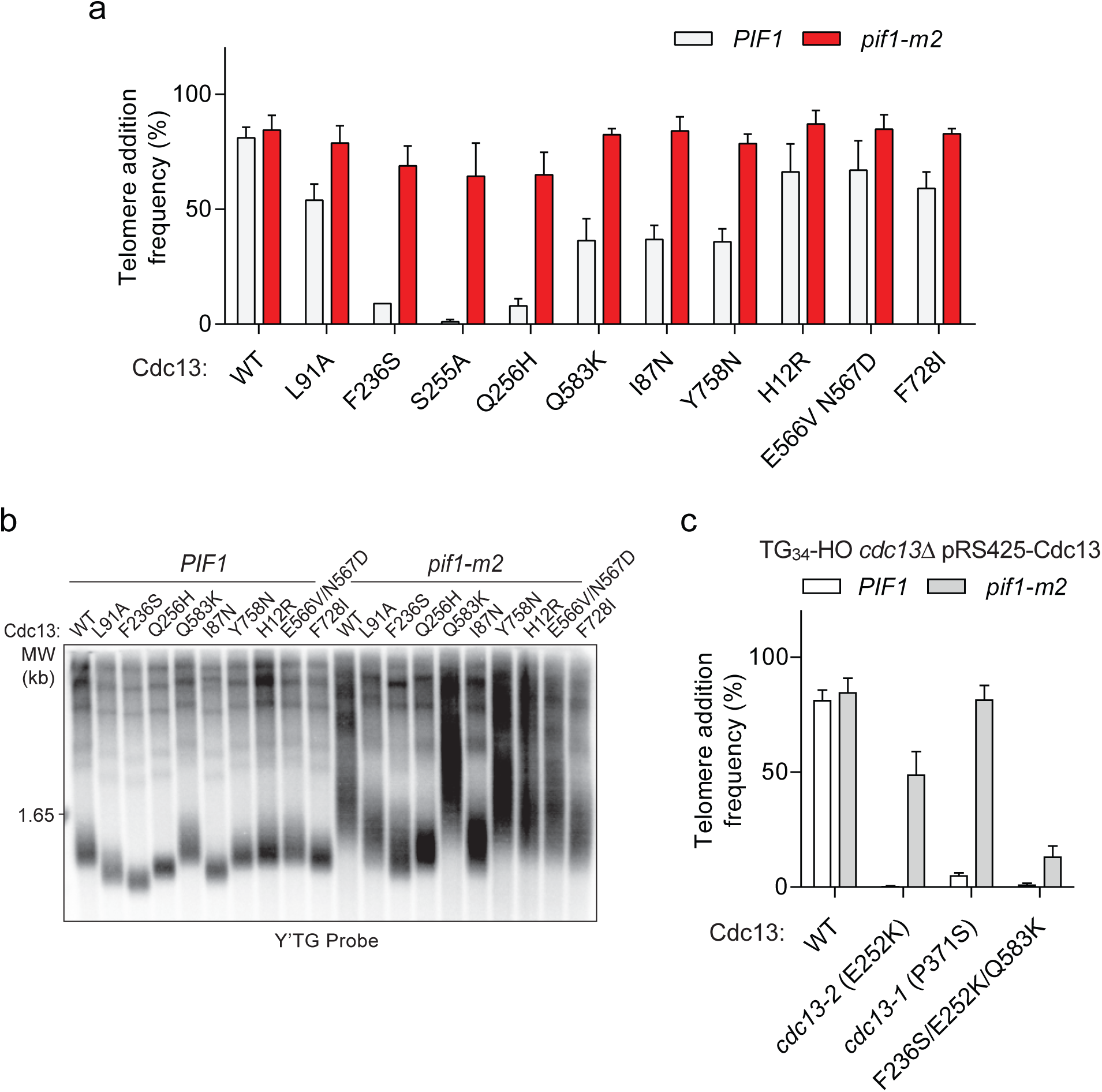
Cdc13 mutations that sensitize the TG_34_ end to Pif1 activity. **a**, Telomere addition frequency at the TG_34_ DNA end in *PIF1* and *pif1-m2* cells in a *cdc13Δ* background expressing wild-type or mutated Cdc13 from pRS425. Data represents the mean ± s.d. from n=3 independent experiments. **b**, Southern blot for telomere length of the strains examined in panel **a**. Cells were passaged for approximately 75 generations before genomic DNA extraction. A Y’TG probe was used to label telomere sequences. **c**, Telomere addition frequency at the TG_34_ DNA end in *PIF1* and *pif1-m2* cells in a *cdc13Δ* background expressing plasmid-borne wild-type or mutated Cdc13. The *cdc13-1* mutant was grown at a permissive temperature of 23°C. Data represents the mean ± s.d. from n=3 independent experiments.

Three powerful single mutations could completely recapitulate the phenotype of the remaining three alleles, namely F236S, S255P, and Q256H (**Figure 7a**). These residues all map to the Cdc13 recruitment domain, suggesting that weakening the association of Cdc13 with telomerase is another means to facilitate Pif1 activity at TG_34_. While the mutation of F236 or Q256 have not been previously reported, the surrounding residues P235, F237, and S255 have been identified to impair telomerase function (Gao et al., 2010; Tseng et al., 2006). Telomere length in several of the *cdc13-sp* alleles was reduced in both wild-type and *pif1-m2* backgrounds, and the severity of the defect generally correlated with the magnitude of the telomere addition phenotype (**Figure 7b**).

The diversity of Cdc13 mutations that sensitize the TG_34_ end to Pif1 suggests that generally disrupting Cdc13 function facilitates Pif1 activity by shifting the balance away from telomere addition. In agreement with this idea, the classic telomerase null *cdc13-2* allele (Lendvay et al., 1996; Nugent et al., 1996) was also sensitive to Pif1, a phenotype that was mirrored in *cdc13-1* mutants grown at permissive temperature (**Figure 7c**); a mutation now known to disrupt OB2 dimerization (Mason et al., 2013). Furthermore, analysis of hits from our screen that decreased telomere addition in both wild-type and *pif1-m2* cells revealed double mutations of critical residues including S255L/Q256R in clone 37, I87T/F236Y in clone 40, and F236S/E252K in clone 48, suggesting that further disrupting Cdc13 function eventually impairs telomere addition even in the absence of *PIF1*. In line with this idea, the F235S/E252K/Q583K triple mutant prevented telomere addition at a TG_34_ end even in *pif1-m2* cells (**Figure 7c**).

### Pif1 does not limit elongation of longer telomeric seeds

A final observation regarding the DSB-telomere transition involves the extension of longer telomeric repeats. While the TG_34_ and TG_82_ substrates are rapidly elongated by telomerase following DNA cleavage, longer repeats such as TG_250_ have been observed by Southern blot to be relatively inert (Negrini et al., 2007). This observation is consistent with the established ability of cells to preferentially elongate short telomeres (Marcand et al., 1999; Teixeira et al., 2004). To further characterize this phenomenon, we generated strains that would yield 96, 119, 142, and 162 bp of telomeric sequence following HO-induced cleavage and followed DNA ends by Southern blot. While minor elongation was detectable at the TG_119_ end, the TG_96_, TG_14_2, and TG_162_ ends were stable within the timescale of the experiment (**Figure 8ab**). The increased extension of TG_119_ compared to the TG_96_ end suggests that specific features of the TG_96_ substrate make it refractory to telomerase and not the length *per se*. A prime candidate for this regulation is the number and the affinity of Rap1 proteins that can bind to the array (Negrini et al., 2007).

**Figure 8.**
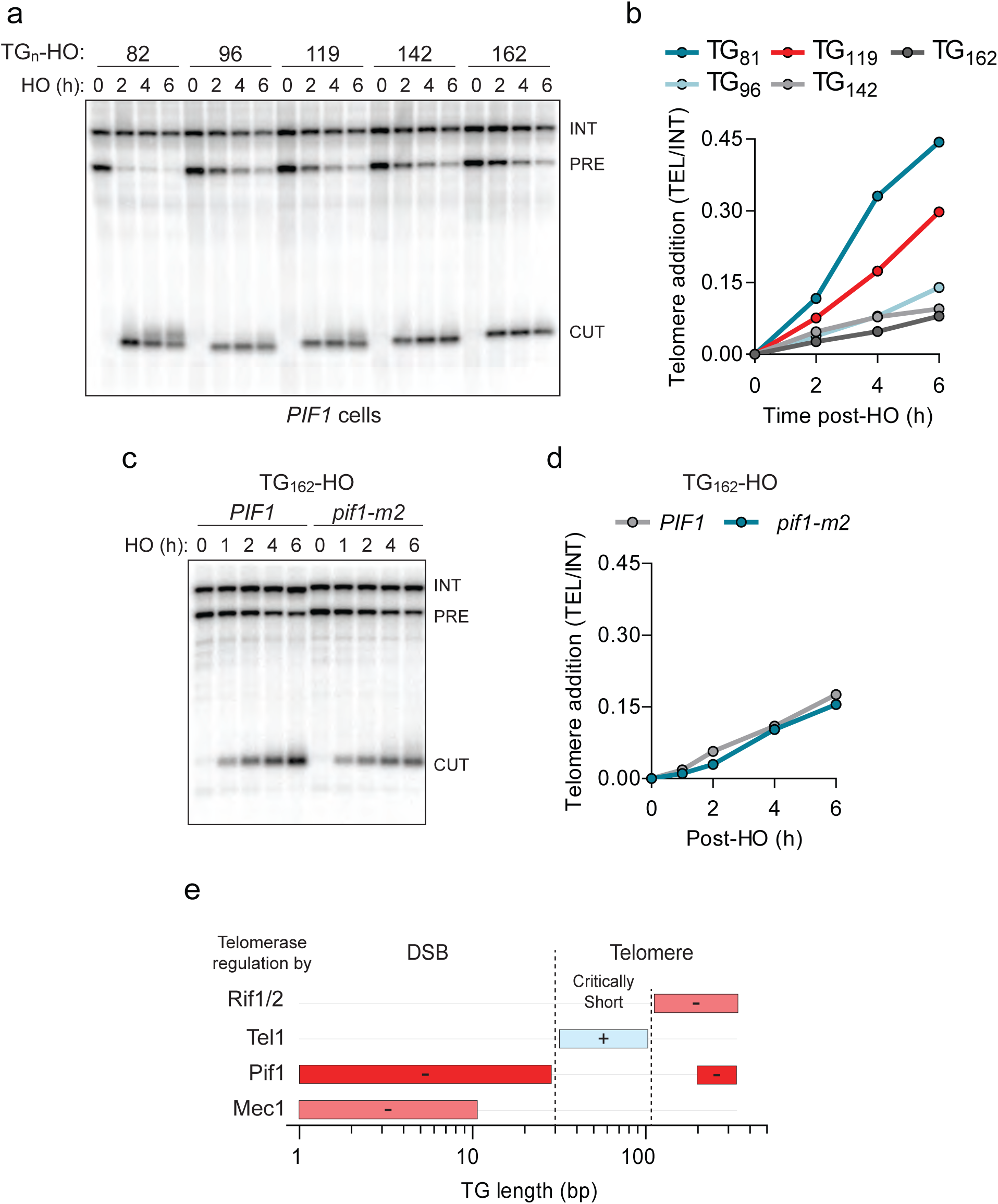
The preferential extension of short telomeres is independent of Pif1. **a**, Southern blot of DNA ends containing 82-162 bp of (TG_1_-3)_n_ sequence following HO induction. A *URA3* probe was used to label the *ura3-52* internal control (INT) and the *URA3* gene adjacent to the TG_n_-HO insert (PRE) that is cleaved by HO endonuclease (CUT). **b**, Quantification of the newly added telomere signal in panel **a** calculated by subtracting the signal prior to HO-induced and normalizing to the internal control. Data represents n=1 experiment. **c**, Southern blot of the TG_162_ DNA end in wild-type (WT) and *pif1-m2* cells following HO induction. **d**, Quantitation of the experiment in panel **c** as described for (**b**). Data represents n=1 experiment. **e**, A model for the length-dependent regulation of telomerase at DNA ends. See the discussion for more detail.

The limited extension of the TG_162_ substrate raised the possibility that Pif1 might counteract telomerase at longer TG repeats, thereby promoting the preferential elongation of short telomeres. This possibility is supported by the recent observation that Pif1 association is increased at long telomeres (Phillips et al., 2015). Contrary to this idea, however, we found that the loss of Pif1 did not increase elongation of the TG_162_ DNA end (**Figure 8cd**) indicating that additional regulatory mechanisms promote extension by telomerase. Together these results suggest that DNA ends containing between 34 and approximately 120 bp of telomeric repeat sequence are recognized as critically short telomeres in budding yeast and are subject to immediate elongation.

## Discussion

The work presented here sheds light on how cells distinguish between DSBs and short telomeres and reveals a striking transition in the fate of DNA ends with regards to the activity of the telomerase inhibitor Pif1. Our findings agree with previous reports demonstrating that linear plasmid substrates containing 41 bp of telomeric repeats are efficiently converted into telomeres (Lustig, 1992) and that a TG_22_ sequence can promote telomere addition (Hirano and Sugimoto, 2007). The discovery here that increased telomere addition is due to the apparent inactivity of Pif1 is a novel insight and future work will be required to fully characterize the molecular switch that occurs at the DSB-telomere transition.

The observed behavior of Pif1 complements several known mechanisms that tightly integrate telomeric sequence length and the regulation of telomerase (**Figure 8e**). The identified activity of Pif1 at telomeric repeats under 34 bp joins a Mec1-dependent mechanism that inhibits Cdc13 binding at repeats under 11 bp (Zhang and Durocher, 2010), highlighting the importance of inhibiting telomerase at DSBs. Conversely, we propose that DNA ends containing telomeric sequences of 34 bp to approximately 120 bp are recognized as critically short telomeres and are preferentially elongated. Tel1 is implicated as a key regulator in this process (Arneric and Lingner, 2007; Chang et al., 2007; Cooley et al., 2014), although the exact phosphorylation targets are unknown (Gao et al., 2010). Finally, the canonical counting mechanism of telomeres is known to limit the extension of long telomeres through the negative regulators Rif1 and Rif2 (Hirano et al., 2009; Levy and Blackburn, 2004; Marcand et al., 1997; McGee et al., 2010). Although our data indicates that Pif1 is not responsible for the initial stability of the TG_162_ end, it has been proposed that Pif1 promotes the preferential elongation of short telomeres by specifically interacting with longer telomeres (Phillips et al., 2015).

In order to maintain genome stability, one might imagine that the length of telomeric repeat sequence necessary to overcome Pif1 should be greater than any natural sequence occurring within in the genome. Any longer sequences should therefore be prone to conversion into new telomeres and might be under negative selection during evolution due to the loss of genetic material. Consistent with this idea, the two longest (TG_1_-3)_n_ repeats in the correct orientation outside of telomeric regions in budding yeast include a 35 bp sequence on Chr VII, and a 31 bp sequence on Chr VI (Mangahas et al., 2001).

Our investigation into the molecular trigger of the DSB-telomere transition points to a key role for the DNA binding protein Cdc13. This conclusion is supported by work revealing that microsatellite repeats containing Cdc13 binding sites stimulate telomere addition (Piazza et al., 2012), and recently that a hotspot on Chr V also promotes Cdc13 binding and telomere addition (Obodo et al., 2016). Furthermore, the tethering of Cdc13, but not Rap1, to this site was shown to be sufficient for the formation of new telomeres (Obodo et al., 2016).

The ability of the Cdc13 OB1 domain to dimerize and bind DNA provides an attractive solution to the DSB-telomere transition; however, our results clearly indicate that sensitivity to Pif1 is not unique to any one domain and can result from a variety of mutations throughout Cdc13, most notably in the recruitment domain. Weakening the ability of Cdc13 to recruit telomerase provides a satisfying explanation for sensitivity of the TG_34_ end to Pif1, but is unable to explain why the TG_34_ end is resistant to Pif1 in the first place, especially as fusing telomerase to Cdc13 was unable to overwhelm Pif1 at the TG_18_ substrate. Interestingly, the mammalian CST complex can bind single-stranded telomeric DNA 32 bp and longer (Miyake et al., 2009) suggesting that Cdc13 in combination with Stn1 and Ten1 may also possess unique binding properties.

One key unresolved issue is the mechanism by which Pif1 inhibits telomerase on either side of the DSB-telomere transition and our results with the *pif1-4A* and *-4D* alleles suggest that these activities may be distinct. It is clear that Pif1 can remove telomerase RNA from telomeres (Boulé et al., 2005; Li et al., 2014), but genetic data reveals that Pif1 also has telomerase-independent activity as *PIF1* loss increases the growth of *cdc13-1 tlc1Δ* cells (Dewar and Lydall, 2010). One potential activity for Pif1 at DSBs is through the promotion of DNA end resection, first observed in *cdc13-1* mutants (Dewar and Lydall, 2010). Consistent with this possibility, end resection impairs telomere addition, and new telomeres are added closer to DSB sites in *pif1-m2* cells (Chung et al., 2010). This model therefore predicts that the TG_18_ end may be resected with the help of Pif1, but that resection is blocked at the TG_34_ end, thus providing a satisfying explanation as to why tethering telomerase to the TG_18_ end did not increase telomere addition. In line with this prediction, a TG_22_ end was previously observed to partially suppress DNA end resection compared to a TG_11_ substrate (Hirano and Sugimoto, 2007).

In conclusion, using Pif1 as a cellular indicator for the DNA-end fate decision reveals a striking threshold that recapitulates several properties of DSBs and telomeres. We propose that the TG_34_ DNA end, approximately ten percent of a healthy budding yeast telomere, is interpreted by the cell as a minimal telomere and future work will be required to characterize its unique properties.

## Acknowledgements

We are grateful to Rachel Szilard and members of the Durocher laboratory for critical reading of the manuscript. JS is supported by a CIHR Doctoral award. DD is the Thomas Kierans Chair in Mechanisms of Cancer Development and a Canada Research Chair (Tier 1) in the Molecular Mechanisms of Genome Integrity. Work in the DD laboratory was supported by CIHR grant MOP106576 and a Grant-in-Aid from the Krembil Foundation.

## Methods

### Yeast strain construction and growth

Strains were constructed by standard allele replacement, PCR-mediated gene deletion or epitope-tagging methods, or via transformations of the indicated plasmids. The desired mutations were selected by prototrophy or drug selection and verified by PCR or sequencing. Standard yeast media and growth conditions were used. Cells were grown in supplemented minimal medium (SD: 2 g/L amino acids dropout mix, 5 g/L ammonium sulfate, 1.7 g/L yeast nitrogen base without amino acids or ammonium sulfate) or in rich medium (XY: 20 g/L bactopeptone, 10 g/L yeast extract, 0.1 g/L adenine, 0.2 g/L tryptophan), containing 2% glucose, 2% raffinose or 3% galactose as indicated.

Telomeric repeats were cloned into the pVII-L plasmid which features an HO endonuclease cut site, a *URA3* selection marker, and homology arms for integration at the *ADH4* locus (Gottschling et al., 1990). Longer telomeric repeats were assembled using commercial gene synthesis (Mr. Gene) while Quikchange mutagenesis (Stratagene) was performed for further manipulation of repeat sequences. Insertions and deletions of up to 30 bp of TG repeats were robustly obtained in a single round of mutagenesis. Quikchange mediated shortening of a large TG_250_ sequence also yielded a wide range of shorter repeats. All repeats were verified by DNA sequencing before integration.

The TG_82_-HO cassette on Chr VII was replaced by integrating SalI and EcoRI-digested pVII-L plasmids and selecting for colonies on SD-URA. Single integration of the plasmid and HO cleavage at the locus was confirmed by Southern blot. Telomere addition strains were constructed in a *rad52Δ* background with a covering pRS414-Rad52 plasmid to facilitate genome manipulation through homologous recombination. Strains were cured of the pRS414-Rad52 plasmid by random loss in non-selective media and colonies were screened by replica-plating to SD-trp.

Pif1 mutations were generated by Quikchange mutagenesis (Stratagene) on a pAUR101-*pif1-m1* nuclear specific construct and integrated at the *AUR1* locus in *pif1-m2* cells. The *est2-up34* mutation was generated by pop-in/pop-out gene replacement.

### Telomere addition assays

Telomere addition assays were performed as previously described (Zhang and Durocher, 2010). Briefly, yeast cultures were grown overnight in XY + glucose to log phase and subcultured into XY + raffinose (2%) for overnight growth to a density of 2.5-7.5×10^6^ cells/mL. Nocodazole (Sigma Aldrich) was added at 15 µg/mL for 2h to synchronize cells in G2/M before addition of galactose to induce HO endonuclease expression. Cells were plated on XY + glucose plates before the addition of galactose and 4 h after galactose addition, and grown for 2 days. The total number of colonies were counted, following which colonies were replica-plated to media containing α-aminoadipic acid (α-AA) to identify cells that had lost the distal *LYS2* gene on Chr VII. Frequency of telomere addition was calculated as the percent of post-galactose surviving colonies that were α-AA resistant. An alternative calculation, (α-AA resistant colonies/ (pre-galactose colonies - α-AA sensitive colonies)), revealed the same threshold of Pif1 activity between the TG_18_ and TG_34_ ends, but with increased variability between experiments.

### Genomic DNA extraction

Genomic DNA was isolated using a phenol-chloroform extraction protocol. Briefly, overnight cultures of cells were grown to saturation, pelleted, and resuspended with 200 µL ‘Smash & Grab’ lysis buffer (10 mM Tris-Cl, pH 8.0, 1 mM EDTA, 100 mM NaCl, 1% SDS, 2% Triton X-100). 200 µL of glass beads (Sigma Aldrich, 400-600 µm diameter) were added along with 200 µL phenol-chloroform (1:1). Cells were lysed by vortexing for 5 min before addition of 200 µL TE buffer (10 mM Tris-Cl pH 8, 1 mM EDTA). Samples were centrifuged at 4°C and DNA from the upper layer precipitated with the addition of 1 mL ice-cold 100% ethanol and centrifuged at 4°C. The DNA pellets were resuspended in 200 µL TE with 300 µg RNAse A (Sigma) and incubated at 37°C for 30 min. DNA was again precipitated with the addition of 1 mL ice-cold 100% ethanol and 10 µL of 4M ammonium acetate, centrifuged, dried, and resuspended in TE.

### Southern blots for telomere addition and length

Fifteen micrograms of genomic DNA were digested overnight with SpeI (for TG_82_ strains) or EcoRV (for all other TG repeat lengths). Digested DNA was run on a 1% agarose gel in 0.5X TBE buffer (45 mM Tris-borate, 1 mM EDTA) at 100V for 6 hours, denatured in the gel for 30 min with 0.5 M NaOH and 1.5 mM NaCl, and neutralized for 30 min with 1.5 M NaCl and 0.5 M Tris-Cl pH 7.5. DNA was transferred to Hybond N+ membrane (GE Healthcare Life Sciences) using overnight capillary flow and 10X SSC buffer (1.5 M NaCl, 150 mM sodium citrate, pH 7). Membranes were UV-crosslinked (Stratalinker 1800, Stratagene) and blocked at 65°C with Church hybridization buffer (250 mM NaPO4 pH 7.2, 1 mM EDTA, 7% SDS). Radiolabelled probes complementary to the *ADE2* (for TG_82_ strains) or *URA3 gene* (for all other TG repeat lengths) were generated from purified PCR products using the Prime-It Random labelling kit (Stratagene) and α^32^-dCTP. Membranes were probed overnight, washed three times with 65°C Church hybridization buffer and exposed overnight with a phosphor screen (GE Healthcare Life Science) before imaging on a Storm or Typhoon FLA 9000 imager (GE Healthcare Life Sciences). Quantification of the added telomere signal (above CUT band) was performed in ImageQuant GE Healthcare Life Sciences) by subtracting the background signal before HO induction followed by normalization to the internal loading control (INT). Telomere length analysis was performed by digesting genomic DNA with XhoI and probing with a Y’-TG probe generated from the pYT14 plasmid (Shampay et al., 1984).

### PCR mutagenesis screens

Mutant alleles were generated by error-prone PCR using Taq polymerase (New England Biolabs) and 0.25 mM MnCl, and purified using spin columns (Qiagen). The Pif1 mutagenesis screen was performed in TG_82_ *pif1-m2* cells co-transformed with gapped pRS416-*pif1-m1* and purified inserts. Cells harbouring repaired plasmids were selected on SD-ura. The Cdc13 mutagenesis screen was performed in TG_34_ *cdc13Δ* cells containing a covering YEp-*CDC13-URA3* plasmid and co-transformed with gapped pRS425-*CDC13* plasmid and PCR inserts. Cells harbouring repaired plasmids were selected on SD-ura before replica-plating to 5-fluoroorotic acid (5-FOA) to remove the covering plasmid. Mutant *cdc13* alleles that are defective in capping should be inviable at this step. Colonies from both screens were patched onto raffinose plates and grown for 2 days before replica plating to galactose plates for 4 hours, and finally to α-AA plates after reducing cell density by first replicating plating to a blank agar plate. Plasmids were rescued using a phenol-chloroform extraction and transformed into *Escherichia coli*. Plasmids were sequenced to identify mutations and retransformed into the parental yeast strain to confirm that the phenotype resulted from the plasmid mutation.

### Statistics

All statistical analysis was performed with GraphPad Prism v5.02 (GraphPad Software) using one-way analysis of variance (ANOVA) followed by Bonferroni post-hoc analysis.

---

**Supplementary Figure 1.**
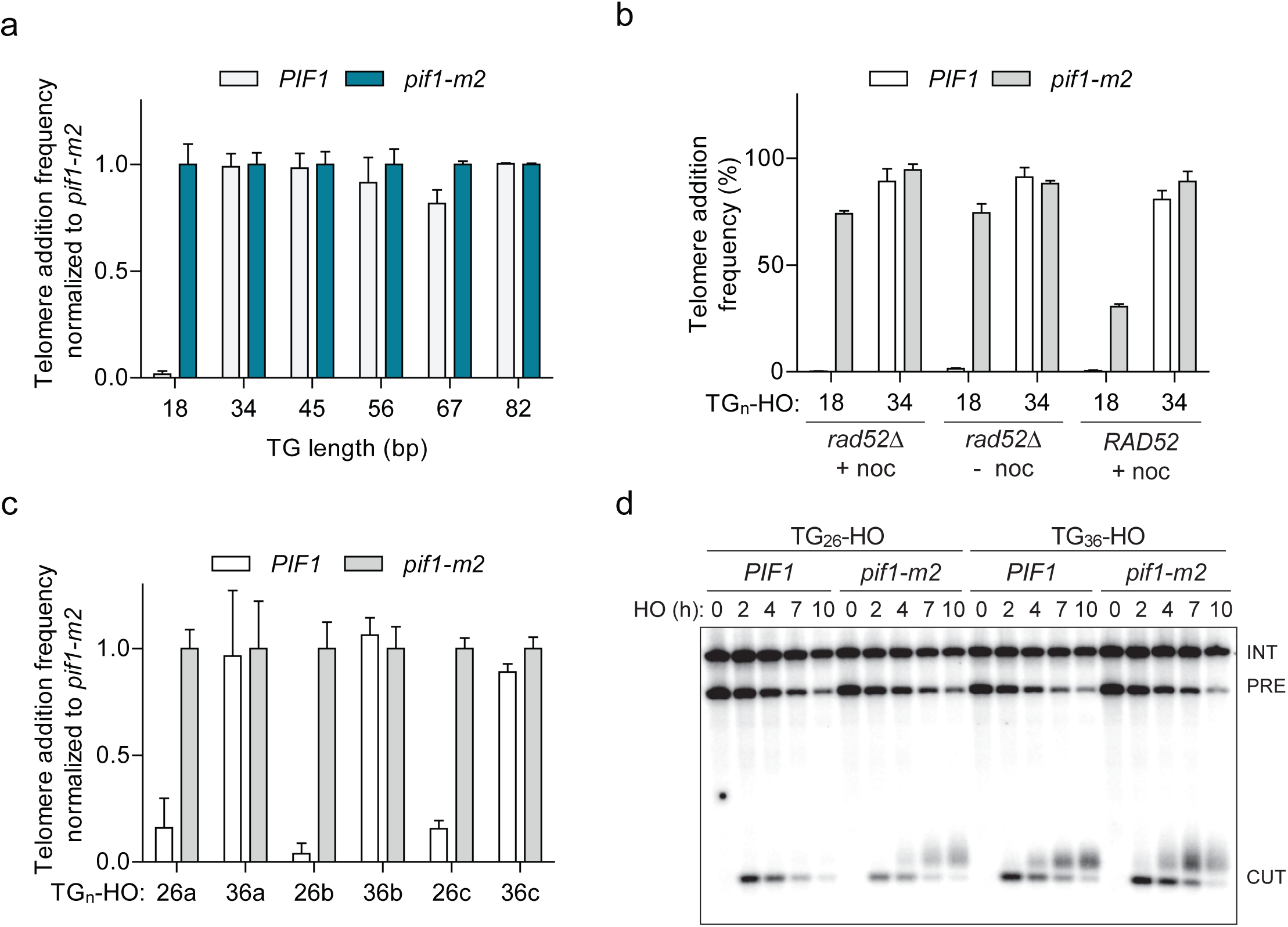
Characterizing a threshold of Pif1 activity at DNA ends. **a**, Telomere addition frequency normalized to *pif1-m2* cells at DNA ends containing 18-82 bp of (TG_1_-3)_n_ sequence. Data represents the mean ± s.d. from n=3 independent experiments. **b**, Telomere addition frequency in *rad52Δ* cells synchronized with 15 µg/mL nocodazole for 2 h (+ noc), asynchronously growing cells (- noc), and in cells harbouring a pRS415-*RAD52* plasmid. Data represents the mean ± s.d. from n=3 independent experiments. **c**, Telomere addition frequency normalized to *pif1-m2* cells at DNA ends containing 26 bp or 36 bp versions of three different natural telomeric (TG_1_-3)_n_ sequences. Data represents the mean ± s.d. from n=3 independent experiments. **d**, Southern blot of DNA ends containing the TG_26b_ and TG_36b_ ends in wild-type (WT) and *pif1-m2* cells following HO induction. A *URA3* probe was used to label the *ura3-52* internal control (INT) and the *URA3* gene adjacent to the TGn-HO insert (PRE) which is cleaved by HO endonuclease (CUT).

**Supplementary Figure 2.**
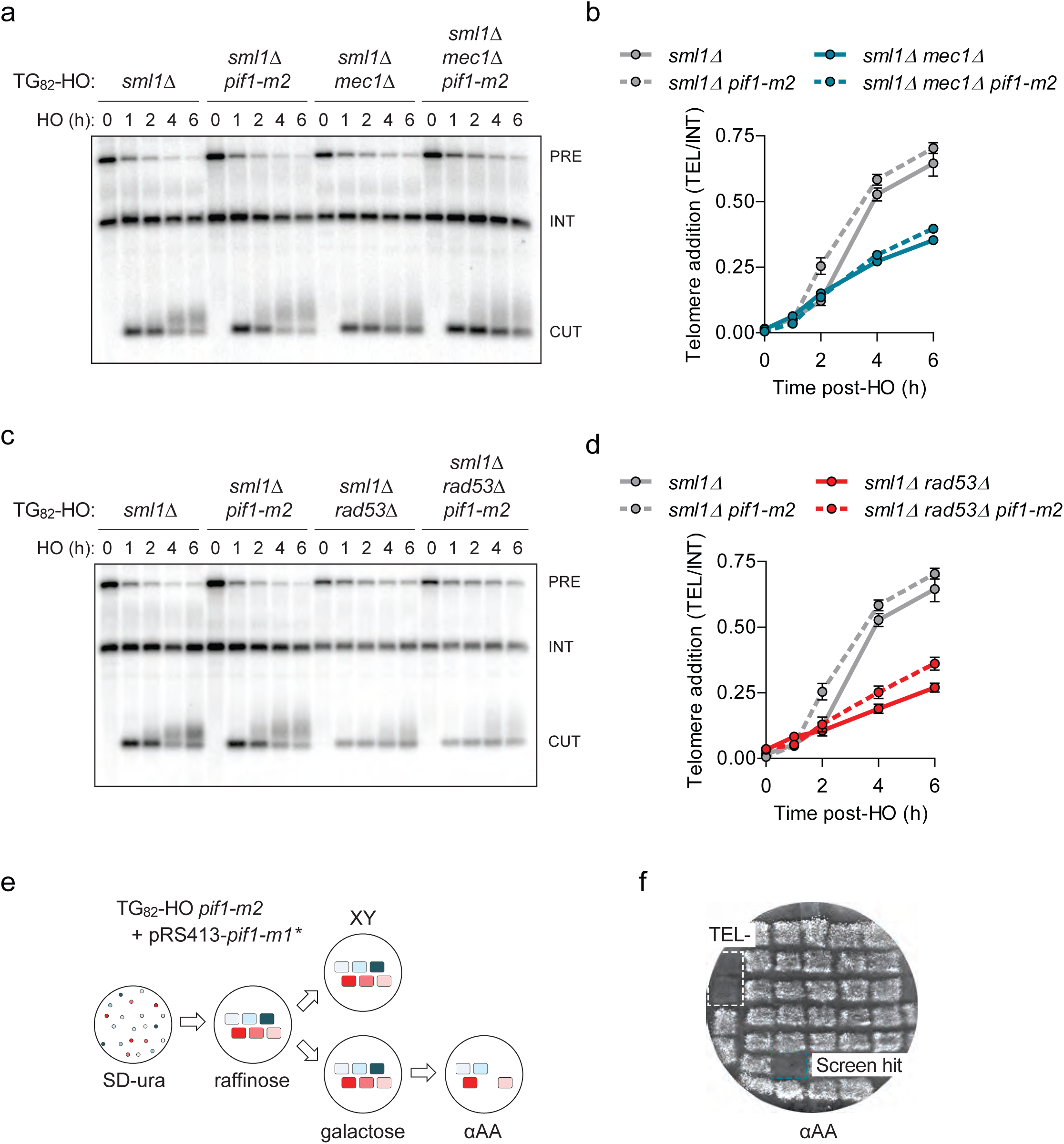
Loss of *MEC1* or *RAD53* does not affect Pif1 at short telomeres. **a-d**, Southern blot of the TG_82_ DNA end following HO induction in *sml1Δ* and *sml1Δ pif1-m2* cells combined with the deletion of *MEC1* (**a**) or *RAD53* (**c**). *SML1* was deleted to suppress the lethality of *mec1Δ* and *rad53Δ.* An *ADE2* probe was used to label the *ade2Δ1* internal control (INT) and the *ADE2* gene adjacent to the TGn-HO insert (PRE) which is cleaved by HO endonuclease (CUT). Quantification of the newly added telomere signal in *mec1Δ* cells (**b**), from n=1 experiment, and *rad53Δ* cells (**d**), from n=2 independent experiments. Data represents the mean ± s.d. **e**, Schematic of a screen in TG_82_ *pif1-m2* cells using a plate-based genetic assay for telomere addition. Cells harbouring repaired mutant *pif1-m1* plasmids were selected on SD-ura before DSB induction by growth for 4 h on galactose medium. A blank agar plate was used to reduce cell number before final selection on α-AA. Unless other indicated, cells were grown on the indicated media for 2-3 days at 30°C. Approximately 2500 Pif1 mutants were screened with an average mutation rate of 0.008 nucleotides/position. **f**, Example plate from the screen. Pif1 mutants that prevent telomere addition are identified by the inability to grow on αAA media (blue box). The TG_18_-HO strain was used as a control since telomere addition is normally inhibited by Pif1 at this substrate (white box).

**Supplementary Table 1:**
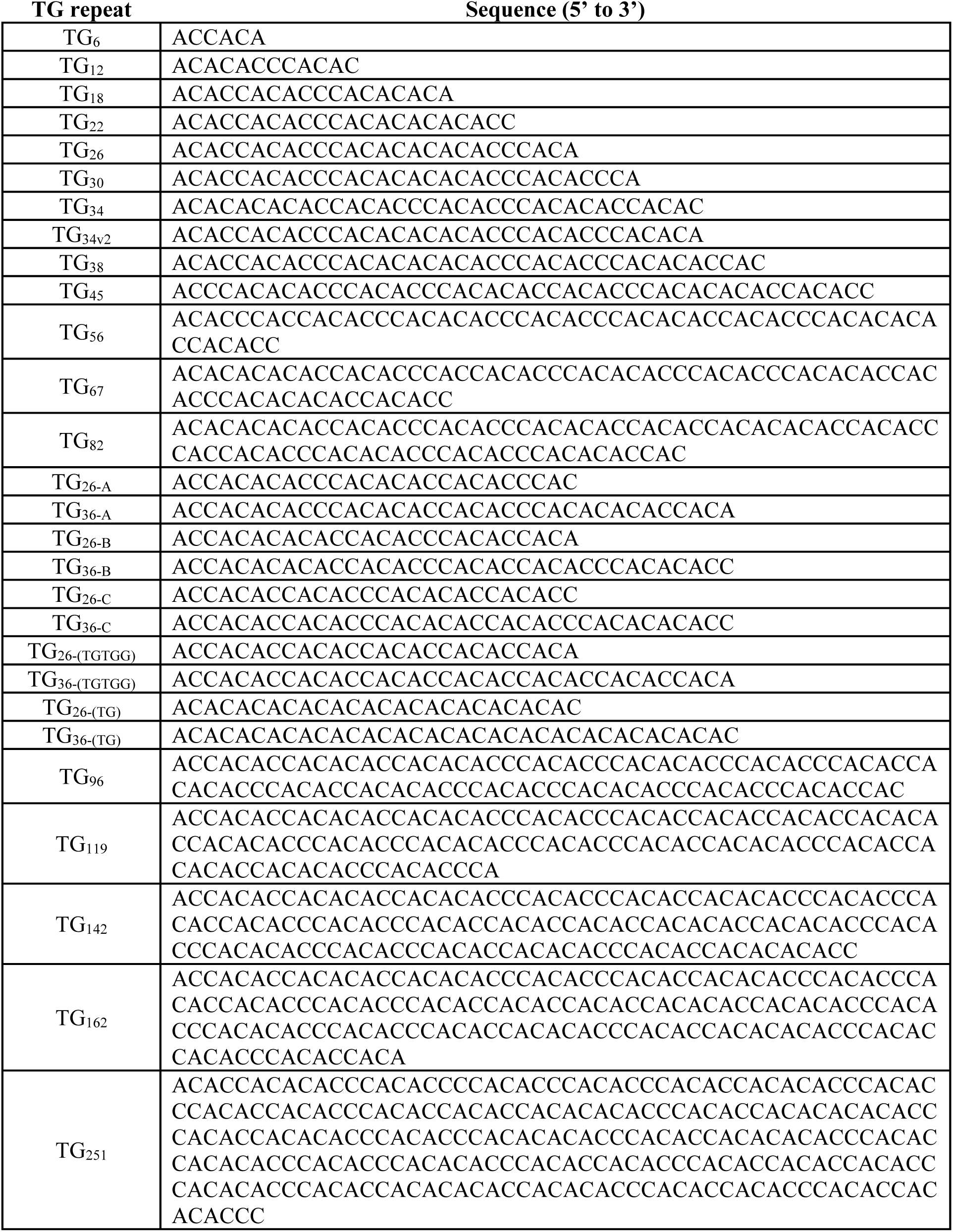
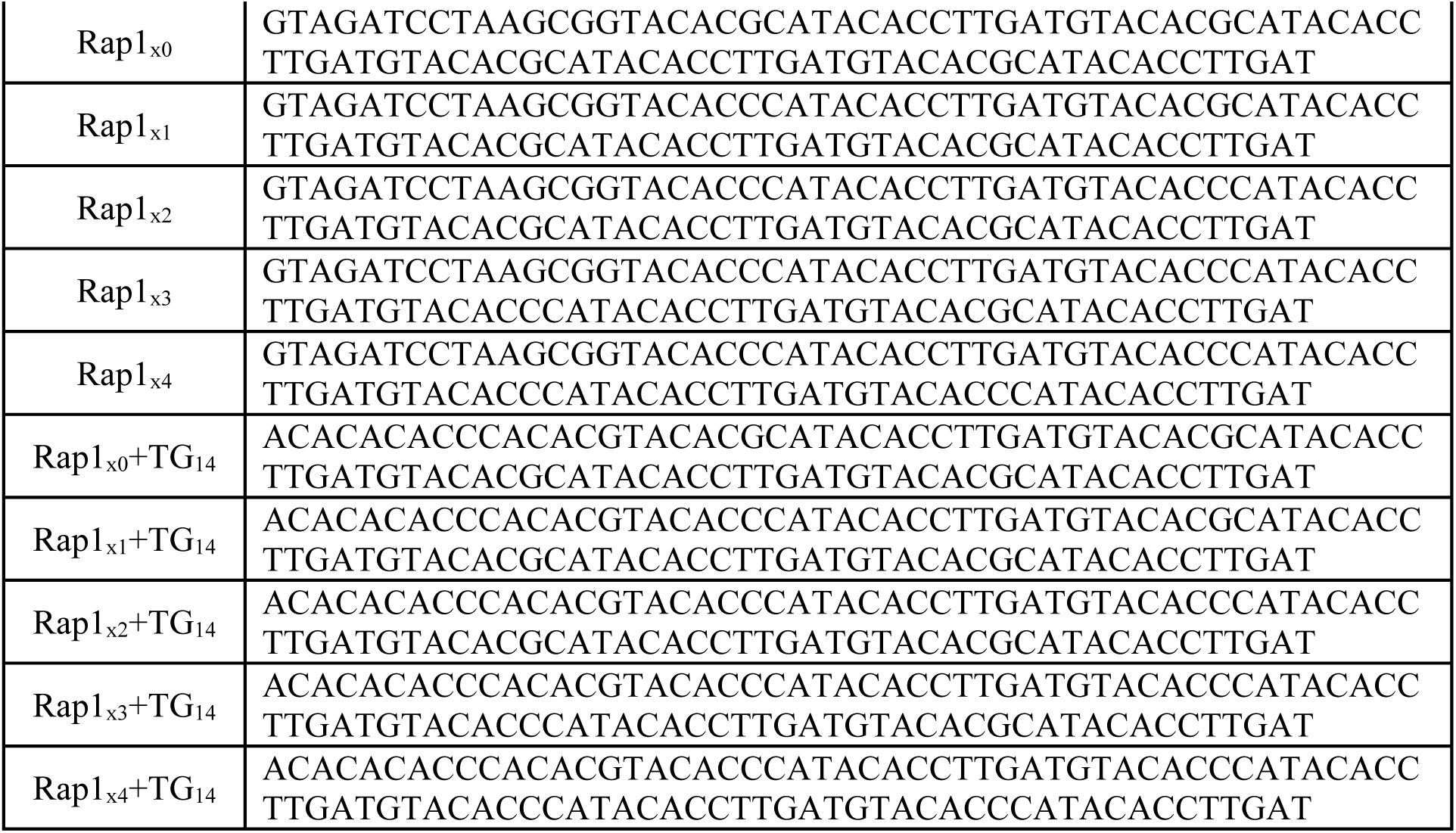
Sequences of DNA ends

**Supplementary Table 2:** Yeast strains

| Strain Number | Strain Background | Genotype | Source |
| --- | --- | --- | --- |
| DDY2458 | S288C | *Mata-inc ura3-52 lys2-801 ade2-101 ochre trp1-Δ63 his3-Δ200 leu2-Δ1::GAL1:HO-LEU2 rad52::HIS pRAD52-TRP VII-L-ADE2-TG82-HOcs-LYS2* | (Zhang & Durocher, 2010) |
| DDY2476 | S288C | *Mata-inc ura3-52 lys2-801 ade2-101 ochre trp1-Δ63 his3-Δ200 leu2-Δ1::GAL1:HO-LEU2 rad52::HIS pRAD52-TRP VII-L-ADE2-TG82-HOcs-LYS2 pif1-m2* | (Zhang & Durocher, 2010) |
| DDY3254 | S288C | *Mata-inc ura3-52 lys2-801 ade2-101 ochre trp1-Δ63 his3-Δ200 leu2-Δ1::GAL1:HO-LEU2 rad52::HIS VII-L::URA3-TG18-HOcs-LYS2* | This study |
| DDY3203 | S288C | *Mata-inc ura3-52 lys2-801 ade2-101 ochre trp1-Δ63 his3-Δ200 leu2-Δ1::GAL1:HO-LEU2 rad52::HIS VII-L::URA3-TG18-HOcs-LYS2 pif1-m2* | This study |
| DDY3376 | S288C | *Mata-inc ura3-52 lys2-801 ade2-101 ochre trp1-Δ63 his3-Δ200 leu2-Δ1::GAL1:HO-LEU2 rad52::HIS VII-L::URA3-TG34-HOcs-LYS2* | This study |
| DDY2986 | S288C | *Mata-inc ura3-52 lys2-801 ade2-101 ochre trp1-Δ63 his3-Δ200 leu2-Δ1::GAL1:HO-LEU2 rad52::HIS VII-L::URA3-TG34-HOcs-LYS2 pif1-m2* | This study |
| DDY2985 | S288C | *Mata-inc ura3-52 lys2-801 ade2-101 ochre trp1-Δ63 his3-Δ200 leu2-Δ1::GAL1:HO-LEU2 rad52::HIS VII-L::URA3-TG45-HOcs-LYS2* | This study |
| DDY2987 | S288C | *Mata-inc ura3-52 lys2-801 ade2-101 ochre trp1-Δ63 his3-Δ200 leu2-Δ1::GAL1:HO-LEU2 rad52::HIS VII-L::URA3-TG45-HOcs-LYS2 pif1-m2* | This study |
| DDY3129 | S288C | *Mata-inc ura3-52 lys2-801 ade2-101 ochre trp1-Δ63 his3-Δ200 leu2-Δ1::GAL1:HO-LEU2 rad52::HIS VII-L::URA3-TG56-HOcs-LYS2* | This study |
| DDY3130 | S288C | *Mata-inc ura3-52 lys2-801 ade2-101 ochre trp1-Δ63 his3-Δ200 leu2-Δ1::GAL1:HO-LEU2 rad52::HIS VII-L::URA3-TG56-HOcs-LYS2 pif1-m2* | This study |
| DDY2988 | S288C | *Mata-inc ura3-52 lys2-801 ade2-101 ochre trp1-Δ63 his3-Δ200 leu2-Δ1::GAL1:HO-LEU2 rad52::HIS VII-L::URA3-TG67-HOcs-LYS2* | This study |
| DDY2990 | S288C | *Mata-inc ura3-52 lys2-801 ade2-101 ochre trp1-Δ63 his3-Δ200 leu2-Δ1::GAL1:HO-LEU2 rad52::HIS VII-L::URA3-TG67-HOcs-LYS2 pif1-m2* | This study |
| DDY2472 | S288C | *Mata-inc ura3-52 lys2-801 ade2-101 ochre trp1-Δ63 his3-Δ200 leu2-Δ1::GAL1:HO-LEU2 rad52::HIS VII-L-ADE2-TG82-HOcs-LYS2* | This study |
| DDY2556 | S288C | *Mata-inc ura3-52 lys2-801 ade2-101 ochre trp1-Δ63 his3-Δ200 leu2-Δ1::GAL1:HO-LEU2 rad52::HIS VII-L-ADE2-TG82-HOcs-LYS2 -m2* | This study |
| DDY3204 | S288C | *Mata-inc ura3-52 lys2-801 ade2-101 ochre trp1-Δ63 his3-Δ200 leu2-Δ1::GAL1:HO-LEU2 rad52::HIS VII-L::URA3-TG22-HOcs-LYS2* | This study |
| DDY3205 | S288C | *Mata-inc ura3-52 lys2-801 ade2-101 ochre trp1-Δ63 his3-Δ200 leu2-Δ1::GAL1:HO-LEU2 rad52::HIS VII-L::URA3-TG22-HOcs-LYS2 pif1-m2* | This study |
| DDY3206 | S288C | *Mata-inc ura3-52 lys2-801 ade2-101 ochre trp1-Δ63 his3-Δ200 leu2-Δ1::GAL1:HO-LEU2 rad52::HIS VII-L::URA3-TG26-HOcs-LYS2* | This study |
| DDY3207 | S288C | *Mata-inc ura3-52 lys2-801 ade2-101 ochre trp1-Δ63 his3-Δ200 leu2-Δ1::GAL1:HO-LEU2 rad52::HIS VII-L::URA3-TG26-HOcs-LYS2 pif1-m2* | This study |
| DDY3208 | S288C | *Mata-inc ura3-52 lys2-801 ade2-101 ochre trp1-Δ63 his3-Δ200 leu2-Δ1::GAL1:HO-LEU2 rad52::HIS VII-L::URA3-TG30-HOcs-LYS2* | This study |
| DDY3209 | S288C | *Mata-inc ura3-52 lys2-801 ade2-101 ochre trp1-Δ63 his3-Δ200 leu2-Δ1::GAL1:HO-LEU2 rad52::HIS VII-L::URA3-TG30-HOcs-LYS2 pif1-m2* | This study |
| DDY3210 | S288C | *Mata-inc ura3-52 lys2-801 ade2-101 ochre trp1-Δ63 his3-Δ200 leu2-Δ1::GAL1:HO-LEU2 rad52::HIS VII-L::URA3-TG34-HOcs-LYS2* | This study |
| DDY3211 | S288C | *Mata-inc ura3-52 lys2-801 ade2-101 ochre trp1-Δ63 his3-Δ200 leu2-Δ1::GAL1:HO-LEU2 rad52::HIS VII-L::URA3-TG34-HOcs-LYS2 pif1-m2* | This study |
| DDY3404 | S288C | *Mata-inc ura3-52 lys2-801 ade2-101 ochre trp1-Δ63 his3-Δ200 leu2-Δ1::GAL1:HO-LEU2 rad52::HIS VII-L::URA3-TG38-HOcs-LYS2* | This study |
| DDY3406 | S288C | *Mata-inc ura3-52 lys2-801 ade2-101 ochre trp1-Δ63 his3-Δ200 leu2-Δ1::GAL1:HO-LEU2 rad52::HIS VII-L::URA3-TG38-HOcs-LYS2 pif1-m2* | This study |
| DDY3275 | S288C | *Mata-inc ura3-52 lys2-801 ade2-101 ochre trp1-Δ63 his3-Δ200 leu2-Δ1::GAL1:HO-LEU2 rad52::HIS VII-L::URA3-TG26a-HOcs-LYS2* | This study |
| DDY3276 | S288C | *Mata-inc ura3-52 lys2-801 ade2-101 ochre trp1-Δ63 his3-Δ200 leu2-Δ1::GAL1:HO-LEU2 rad52::HIS VII-L::URA3-TG26a-HOcs-LYS2 pif1-m2* | This study |
| DDY3277 | S288C | *Mata-inc ura3-52 lys2-801 ade2-101 ochre trp1-Δ63 his3-Δ200 leu2-Δ1::GAL1:HO-LEU2 rad52::HIS VII-L::URA3-TG36a-HOcs-LYS2* | This study |
| DDY3278 | S288C | *Mata-inc ura3-52 lys2-801 ade2-101 ochre trp1-Δ63 his3-Δ200 leu2-Δ1::GAL1:HO-LEU2 rad52::HIS VII-L::URA3-TG36a-HOcs-LYS2 pif1-m2* | This study |
| DDY3279 | S288C | *Mata-inc ura3-52 lys2-801 ade2-101 ochre trp1-Δ63 his3-Δ200 leu2-Δ1::GAL1:HO-LEU2 rad52::HIS VII-L::URA3-TG26b-HOcs-LYS2* | This study |
| DDY3280 | S288C | *Mata-inc ura3-52 lys2-801 ade2-101 ochre trp1-Δ63 his3-Δ200 leu2-Δ1::GAL1:HO-LEU2 rad52::HIS VII-L::URA3-TG26b-HOcs-LYS2 pif1-m2* | This study |
| DDY3281 | S288C | *Mata-inc ura3-52 lys2-801 ade2-101 ochre trp1-Δ63 his3-Δ200 leu2-Δ1::GAL1:HO-LEU2 rad52::HIS VII-L::URA3-TG36b-HOcs-LYS2* | This study |
| DDY3282 | S288C | *Mata-inc ura3-52 lys2-801 ade2-101 ochre trp1-Δ63 his3-Δ200 leu2-Δ1::GAL1:HO-LEU2 rad52::HIS VII-L::URA3-TG36b-HOcs-LYS2 pif1-m2* | This study |
| DDY3283 | S288C | *Mata-inc ura3-52 lys2-801 ade2-101 ochre trp1-Δ63 his3-Δ200 leu2-Δ1::GAL1:HO-LEU2 rad52::HIS VII-L::URA3-TG26c-HOcs-LYS2* | This study |
| DDY3284 | S288C | *Mata-inc ura3-52 lys2-801 ade2-101 ochre trp1-Δ63 his3-Δ200 leu2-Δ1::GAL1:HO-LEU2 rad52::HIS VII-L::URA3-TG26c-HOcs-LYS2 pif1-m2* | This study |
| DDY3285 | S288C | *Mata-inc ura3-52 lys2-801 ade2-101 ochre trp1-Δ63 his3-Δ200 leu2-Δ1::GAL1:HO-LEU2 rad52::HIS VII-L::URA3-TG36c-HOcs-LYS2* | This study |
| DDY3286 | S288C | *Mata-inc ura3-52 lys2-801 ade2-101 ochre trp1-Δ63 his3-Δ200 leu2-Δ1::GAL1:HO-LEU2 rad52::HIS VII-L::URA3-TG36c-HOcs-LYS2 pif1-m2* | This study |
| DDY3042 | S288C | *Mata-inc ura3-52 lys2-801 ade2-101 ochre trp1-Δ63 his3-Δ200 leu2-Δ1::GAL1:HO-LEU2 rad52::HIS VII-L-ADE2-TG82-HOcs-LYS2 tel1::KANMX* | This study |
| DDY3043 | S288C | *Mata-inc ura3-52 lys2-801 ade2-101 ochre trp1-Δ63 his3-Δ200 leu2-Δ1::GAL1:HO-LEU2 rad52::HIS VII-L-ADE2-TG82-HOcs-LYS2 pif1-m2 tel1::KANMX* | This study |
| DDY3483 | S288C | *Mata-inc ura3-52 lys2-801 ade2-101 ochre trp1-Δ63 his3-Δ200 leu2-Δ1::GAL1:HO-LEU2 rad52::HIS VII-L::URA3-TG18-HOcs-LYS2 tel1::KANMX* | This study |
| DDY3484 | S288C | *Mata-inc ura3-52 lys2-801 ade2-101 ochre trp1-Δ63 his3-Δ200 leu2-Δ1::GAL1:HO-LEU2 rad52::HIS VII-L::URA3-TG18-HOcs-LYS2 pif1-m2 tel1::KANMX* | This study |
| DDY3485 | S288C | *Mata-inc ura3-52 lys2-801 ade2-101 ochre trp1-Δ63 his3-Δ200 leu2-Δ1::GAL1:HO-LEU2 rad52::HIS VII-L::URA3-TG34-HOcs-LYS2 tel1::KANMX* | This study |
| DDY3486 | S288C | *Mata-inc ura3-52 lys2-801 ade2-101 ochre trp1-Δ63 his3-Δ200 leu2-Δ1::GAL1:HO-LEU2 rad52::HIS VII-L::URA3-TG34-HOcs-LYS2 pif1-m2 tel1::KANMX* | This study |
| DDY3234 | S288C | *Mata-inc ura3-52 lys2-801 ade2-101 ochre trp1-Δ63 his3-Δ200 leu2-Δ1::GAL1:HO-LEU2 rad52::HIS VII-L::URA3-TG34-HOcs-LYS2 pif1-m2 AUR1* | This study |
| DDY3236 | S288C | *Mata-inc ura3-52 lys2-801 ade2-101 ochre trp1-Δ63 his3-Δ200 leu2-Δ1::GAL1:HO-LEU2 rad52::HIS VII-L::URA3-TG34-HOcs-LYS2 pif1-m2 AUR1::pif1-m1* | This study |
| DDY3244 | S288C | *Mata-inc ura3-52 lys2-801 ade2-101 ochre trp1-Δ63 his3-Δ200 leu2-Δ1::GAL1:HO-LEU2 rad52::HIS VII-L::URA3-TG34-HOcs-LYS2 pif1-m2 AUR1::pif1-m1(5AQ)* | This study |
| DDY3224 | S288C | *Mata-inc ura3-52 lys2-801 ade2-101 ochre trp1-Δ63 his3-Δ200 leu2-Δ1::GAL1:HO-LEU2 rad52::HIS VII-L::URA3-TG18-HOcs-LYS2 pif1-m2 AUR1* | This study |
| DDY3226 | S288C | *Mata-inc ura3-52 lys2-801 ade2-101 ochre trp1-Δ63 his3-Δ200 leu2-Δ1::GAL1:HO-LEU2 rad52::HIS VII-L::URA3-TG18-HOcs-LYS2 pif1-m2 AUR1::pif1-m1* | This study |
| DDY3230 | S288C | *Mata-inc ura3-52 lys2-801 ade2-101 ochre trp1-Δ63 his3-Δ200 leu2-Δ1::GAL1:HO-LEU2 rad52::HIS VII-L::URA3-TG18-HOcs-LYS2 pif1-m2 AUR1::pif1-m1(4A)* | This study |
| DDY3470 | S288C | *Mata-inc ura3-52 lys2-801 ade2-101 ochre trp1-Δ63 his3-Δ200 leu2-Δ1::GAL1:HO-LEU2 rad52::HIS VII-L::URA3-TG18-HOcs-LYS2 pif1-m2 AUR1::pif1-m1(4D)* | This study |
| DDY3240 | S288C | *Mata-inc ura3-52 lys2-801 ade2-101 ochre trp1-Δ63 his3-Δ200 leu2-Δ1::GAL1:HO-LEU2 rad52::HIS VII-L::URA3-TG34-HOcs-LYS2 pif1-m2 AUR1::pif1-m1(4D)* | This study |
| DDY3141 | S288C | *Mata-inc ura3-52 lys2-801 ade2-101 ochre trp1-Δ63 his3-Δ200 leu2-Δ1::GAL1:HO-LEU2 rad52::HIS VII-L-ADE2-TG82-HOcs-LYS2 sml1::NATMX* | This study |
| DDY3142 | S288C | *Mata-inc ura3-52 lys2-801 ade2-101 ochre trp1-Δ63 his3-Δ200 leu2-Δ1::GAL1:HO-LEU2 rad52::HIS VII-L-ADE2-TG82-HOcs-LYS2 pif1-m2 sml1::NATMX* | This study |
| DDY3144 | S288C | *Mata-inc ura3-52 lys2-801 ade2-101 ochre trp1-Δ63 his3-Δ200 leu2-Δ1::GAL1:HO-LEU2 rad52::HIS VII-L-ADE2-TG82-HOcs-LYS2 sml1:NATMX; mec1::KANMX* | This study |
| DDY3039 | S288C | *Mata-inc ura3-52 lys2-801 ade2-101 ochre trp1-Δ63 his3-Δ200 leu2-Δ1::GAL1:HO-LEU2 rad52::HIS VII-L-ADE2-TG82-HOcs-LYS2 pif1-m2 sml1:NATMX; mec1::KANMX* | This study |
| DDY3146 | S288C | *Mata-inc ura3-52 lys2-801 ade2-101 ochre trp1-Δ63 his3-Δ200 leu2-Δ1::GAL1:HO-LEU2 rad52::HIS VII-L-ADE2-TG82-HOcs-LYS2 sml1::NATMX rad53::KANMX* | This study |
| DDY3041 | S288C | *Mata-inc ura3-52 lys2-801 ade2-101 ochre trp1-Δ63 his3-Δ200 leu2-Δ1::GAL1:HO-LEU2 rad52::HIS VII-L-ADE2-TG82-HOcs-LYS2 pif1-m2 sml1::NATMX rad53::KANMX* | This study |
| DDY3224 | S288C | *Mata-inc ura3-52 lys2-801 ade2-101 ochre trp1-Δ63 his3-Δ200 leu2-Δ1::GAL1:HO-LEU2 rad52::HIS VII-L::URA3-TG18-HOcs-LYS2 pif1-m2 AUR1* | This study |
| DDY3226 | S288C | *Mata-inc ura3-52 lys2-801 ade2-101 ochre trp1-Δ63 his3-Δ200 leu2-Δ1::GAL1:HO-LEU2 rad52::HIS VII-L::URA3-TG18-HOcs-LYS2 pif1-m2 AUR1::pif1-m1* | This study |
| DDY3470 | S288C | *Mata-inc ura3-52 lys2-801 ade2-101 ochre trp1-Δ63 his3-Δ200 leu2-Δ1::GAL1:HO-LEU2 rad52::HIS VII-L::URA3-TG18-HOcs-LYS2 pif1-m2 AUR1::pif1-m1(4D)* | This study |
| DDY3604 | S288C | *Mata-inc ura3-52 lys2-801 ade2-101 ochre trp1-Δ63 his3-Δ200 leu2-Δ1::GAL1:HO-LEU2 rad52::HIS VII-L::TG18-HOcs-LYS2 ura3::HPHMX cdc13::KANMX pRS414-Cdc13-Est1* | This study |
| DDY3605 | S288C | *Mata-inc ura3-52 lys2-801 ade2-101 ochre trp1-Δ63 his3-Δ200 leu2-Δ1::GAL1:HO-LEU2 rad52::HIS VII-L::TG18-HOcs-LYS2 ura3::HPHMX cdc13::KANMX pif1-m2 pRS414-Cdc13-Est1* | This study |
| DDY3606 | S288C | *Mata-inc ura3-52 lys2-801 ade2-101 ochre trp1-Δ63 his3-Δ200 leu2-Δ1::GAL1:HO-LEU2 rad52::HIS VII-L::TG34-HOcs-LYS2 ura3::HPHMX cdc13::KANMX pRS414-Cdc13-Est1* | This study |
| DDY3607 | S288C | *Mata-inc ura3-52 lys2-801 ade2-101 ochre trp1-Δ63 his3-Δ200 leu2-Δ1::GAL1:HO-LEU2 rad52::HIS VII-L::TG34-* | This study |
|  |  | *HOcs-LYS2 ura3::HPHMX cdc13::KANMX pif1-m2 pRS414-Cdc13-Est1* |  |
| DDY3608 | S288C | *Mata-inc ura3-52 lys2-801 ade2-101 ochre trp1-Δ63 his3-Δ200 leu2-Δ1::GAL1:HO-LEU2 rad52::HIS VII-L::TG18-HOcs-LYS2 ura3::HPHMX cdc13::KANMX pRS414-Cdc13-Est2* | This study |
| DDY3609 | S288C | *Mata-inc ura3-52 lys2-801 ade2-101 ochre trp1-Δ63 his3-Δ200 leu2-Δ1::GAL1:HO-LEU2 rad52::HIS VII-L::TG18-HOcs-LYS2 ura3::HPHMX cdc13::KANMX pif1-m2 pRS414-Cdc13-Est2* | This study |
| DDY3610 | S288C | *Mata-inc ura3-52 lys2-801 ade2-101 ochre trp1-Δ63 his3-Δ200 leu2-Δ1::GAL1:HO-LEU2 rad52::HIS VII-L::TG34-HOcs-LYS2 ura3::HPHMX cdc13::KANMX pRS414-Cdc13-Est2* | This study |
| DDY3611 | S288C | *Mata-inc ura3-52 lys2-801 ade2-101 ochre trp1-Δ63 his3-Δ200 leu2-Δ1::GAL1:HO-LEU2 rad52::HIS VII-L::TG34-HOcs-LYS2 ura3::HPHMX cdc13::KANMX pif1-m2 pRS414-Cdc13-Est2* | This study |
| DDY3499 | S288C | *Mata-inc ura3-52 lys2-801 ade2-101 ochre trp1-Δ63 his3-Δ200 leu2-Δ1::GAL1:HO-LEU2 rad52::HIS VII-L::URA3-TG18-HOcs-LYS2 est2-up34* | This study |
| DDY3500 | S288C | *Mata-inc ura3-52 lys2-801 ade2-101 ochre trp1-Δ63 his3-Δ200 leu2-Δ1::GAL1:HO-LEU2 rad52::HIS VII-L::URA3-TG18-HOcs-LYS2 pif1-m2 est2up-34* | This study |
| DDY3501 | S288C | *Mata-inc ura3-52 lys2-801 ade2-101 ochre trp1-Δ63 his3-Δ200 leu2-Δ1::GAL1:HO-LEU2 rad52::HIS VII-L::URA3-TG34-HOcs-LYS2 est2-up34* | This study |
| DDY3502 | S288C | *Mata-inc ura3-52 lys2-801 ade2-101 ochre trp1-Δ63 his3-Δ200 leu2-Δ1::GAL1:HO-LEU2 rad52::HIS VII-L::URA3-TG34-HOcs-LYS2 pif1-m2 est2-up34* | This study |
| DDY3287 | S288C | *Mata-inc ura3-52 lys2-801 ade2-101 ochre trp1-Δ63 his3-Δ200 leu2-Δ1::GAL1:HO-LEU2 rad52::HIS VII-L::URA3-TG26d-HOcs-LYS2* | This study |
| DDY3288 | S288C | *Mata-inc ura3-52 lys2-801 ade2-101 ochre trp1-Δ63 his3-Δ200 leu2-Δ1::GAL1:HO-LEU2 rad52::HIS VII-L::URA3-TG26d-HOcs-LYS2 pif1-m2* | This study |
| DDY3289 | S288C | *Mata-inc ura3-52 lys2-801 ade2-101 ochre trp1-Δ63 his3-Δ200 leu2-Δ1::GAL1:HO-LEU2 rad52::HIS VII-L::URA3-TG36d-HOcs-LYS2* | This study |
| DDY3290 | S288C | *Mata-inc ura3-52 lys2-801 ade2-101 ochre trp1-Δ63 his3-Δ200 leu2-Δ1::GAL1:HO-LEU2 rad52::HIS VII-L::URA3-TG36d-HOcs-LYS2 pif1-m2* | This study |
| DDY3291 | S288C | *Mata-inc ura3-52 lys2-801 ade2-101 ochre trp1-Δ63 his3-Δ200 leu2-Δ1::GAL1:HO-LEU2 rad52::HIS VII-L::URA3-TG26e-HOcs-LYS2* | This study |
| DDY3292 | S288C | *Mata-inc ura3-52 lys2-801 ade2-101 ochre trp1-Δ63 his3-Δ200 leu2-Δ1::GAL1:HO-LEU2 rad52::HIS VII-L::URA3-TG26e-HOcs-LYS2 pif1-m2* | This study |
| DDY3293 | S288C | *Mata-inc ura3-52 lys2-801 ade2-101 ochre trp1-Δ63 his3-Δ200 leu2-Δ1::GAL1:HO-LEU2 rad52::HIS VII-L::URA3-TG36e-HOcs-LYS2* | This study |
| DDY3294 | S288C | *Mata-inc ura3-52 lys2-801 ade2-101 ochre trp1-Δ63 his3-Δ200 leu2-Δ1::GAL1:HO-LEU2 rad52::HIS VII-L::URA3-TG36e-HOcs-LYS2 pif1-m2* | This study |
| DDY3324 | S288C | *Mata-inc ura3-52 lys2-801 ade2-101 ochre trp1-Δ63 his3-Δ200 leu2-Δ1::GAL1:HO-LEU2 rad52::HIS VII-L::URA3-Rap1x0+TG14-HOcs-LYS2* | This study |
| DDY3325 | S288C | *Mata-inc ura3-52 lys2-801 ade2-101 ochre trp1-Δ63 his3-Δ200 leu2-Δ1::GAL1:HO-LEU2 rad52::HIS VII-L::URA3-Rap1x0+TG14-HOcs-LYS2 pif1-m2* | This study |
| DDY3326 | S288C | *Mata-inc ura3-52 lys2-801 ade2-101 ochre trp1-Δ63 his3-Δ200 leu2-Δ1::GAL1:HO-LEU2 rad52::HIS VII-L::URA3-Rap1x1+TG14-HOcs-LYS2* | This study |
| DDY3327 | S288C | *Mata-inc ura3-52 lys2-801 ade2-101 ochre trp1-Δ63 his3-Δ200 leu2-Δ1::GAL1:HO-LEU2 rad52::HIS VII-L::URA3-Rap1x1+TG14-HOcs-LYS2 pif1-m2* | This study |
| DDY3328 | S288C | *Mata-inc ura3-52 lys2-801 ade2-101 ochre trp1-Δ63 his3-Δ200 leu2-Δ1::GAL1:HO-LEU2 rad52::HIS VII-L::URA3-Rap1x2+TG14-HOcs-LYS2* | This study |
| DDY3329 | S288C | *Mata-inc ura3-52 lys2-801 ade2-101 ochre trp1-Δ63 his3-Δ200 leu2-Δ1::GAL1:HO-LEU2 rad52::HIS VII-L::URA3-Rap1x2+TG14-HOcs-LYS2 pif1-m2* | This study |
| DDY3330 | S288C | *Mata-inc ura3-52 lys2-801 ade2-101 ochre trp1-Δ63 his3-Δ200 leu2-Δ1::GAL1:HO-LEU2 rad52::HIS VII-L::URA3-Rap1x3+TG14-HOcs-LYS2* | This study |
| DDY3331 | S288C | *Mata-inc ura3-52 lys2-801 ade2-101 ochre trp1-Δ63 his3-Δ200 leu2-Δ1::GAL1:HO-LEU2 rad52::HIS VII-L::URA3-Rap1x3+TG14-HOcs-LYS2 pif1-m2* | This study |
| DDY3332 | S288C | *Mata-inc ura3-52 lys2-801 ade2-101 ochre trp1-Δ63 his3-Δ200 leu2-Δ1::GAL1:HO-LEU2 rad52::HIS VII-L::URA3-Rap1x4+TG14-HOcs-LYS2* | This study |
| DDY3333 | S288C | *Mata-inc ura3-52 lys2-801 ade2-101 ochre trp1-Δ63 his3-Δ200 leu2-Δ1::GAL1:HO-LEU2 rad52::HIS VII-L::URA3-Rap1x4+TG14-HOcs-LYS2 pif1-m2* | This study |
| DDY3334 | S288C | *Mata-inc ura3-52 lys2-801 ade2-101 ochre trp1-Δ63 his3-Δ200 leu2-Δ1::GAL1:HO-LEU2 rad52::HIS VII-L::URA3-Rap1x0-HOcs-LYS2* | This study |
| DDY3335 | S288C | *Mata-inc ura3-52 lys2-801 ade2-101 ochre trp1-Δ63 his3-Δ200 leu2-Δ1::GAL1:HO-LEU2 rad52::HIS VII-L::URA3-Rap1x0-HOcs-LYS2 pif1-m2* | This study |
| DDY3336 | S288C | *Mata-inc ura3-52 lys2-801 ade2-101 ochre trp1-Δ63 his3-Δ200 leu2-Δ1::GAL1:HO-LEU2 rad52::HIS VII-L::URA3-Rap1x1-HOcs-LYS2* | This study |
| DDY3337 | S288C | *Mata-inc ura3-52 lys2-801 ade2-101 ochre trp1-Δ63 his3-Δ200 leu2-Δ1::GAL1:HO-LEU2 rad52::HIS VII-L::URA3-Rap1x1-HOcs-LYS2 pif1-m2* | This study |
| DDY3338 | S288C | *Mata-inc ura3-52 lys2-801 ade2-101 ochre trp1-Δ63 his3-Δ200 leu2-Δ1::GAL1:HO-LEU2 rad52::HIS VII-L::URA3-Rap1x2-HOcs-LYS2* | This study |
| DDY3339 | S288C | *Mata-inc ura3-52 lys2-801 ade2-101 ochre trp1-Δ63 his3-Δ200 leu2-Δ1::GAL1:HO-LEU2 rad52::HIS VII-L::URA3-Rap1x2-HOcs-LYS2 pif1-m2* | This study |
| DDY3340 | S288C | *Mata-inc ura3-52 lys2-801 ade2-101 ochre trp1-Δ63 his3-Δ200 leu2-Δ1::GAL1:HO-LEU2 rad52::HIS VII-L::URA3-Rap1x3-HOcs-LYS2* | This study |
| DDY3341 | S288C | *Mata-inc ura3-52 lys2-801 ade2-101 ochre trp1-Δ63 his3-Δ200 leu2-Δ1::GAL1:HO-LEU2 rad52::HIS VII-L::URA3-Rap1x3-HOcs-LYS2 pif1-m2* | This study |
| DDY3342 | S288C | *Mata-inc ura3-52 lys2-801 ade2-101 ochre trp1-Δ63 his3-Δ200 leu2-Δ1::GAL1:HO-LEU2 rad52::HIS VII-L::URA3-Rap1x4-HOcs-LYS2* | This study |
| DDY3343 | S288C | *Mata-inc ura3-52 lys2-801 ade2-101 ochre trp1-Δ63 his3-Δ200 leu2-Δ1::GAL1:HO-LEU2 rad52::HIS VII-L::URA3-Rap1x4-HOcs-LYS2 pif1-m2* | This study |
| DDY3475 | S288C | *Mata-inc ura3-52 lys2-801 ade2-101 ochre trp1-Δ63 his3-Δ200 leu2-Δ1::GAL1:HO-LEU2 rad52::HIS VII-L::URA3-TG18-HOcs-LYS2 rif1::KANMX* | This study |
| DDY3476 | S288C | *Mata-inc ura3-52 lys2-801 ade2-101 ochre trp1-Δ63 his3-Δ200 leu2-Δ1::GAL1:HO-LEU2 rad52::HIS VII-L::URA3-TG18-HOcs-LYS2 pif1-m2 rif1::KANMX* | This study |
| DDY3477 | S288C | *Mata-inc ura3-52 lys2-801 ade2-101 ochre trp1-Δ63 his3-Δ200 leu2-Δ1::GAL1:HO-LEU2 rad52::HIS VII-L::URA3-TG34-HOcs-LYS2 rif1::KANMX* | This study |
| DDY3478 | S288C | *Mata-inc ura3-52 lys2-801 ade2-101 ochre trp1-Δ63 his3-Δ200 leu2-Δ1::GAL1:HO-LEU2 rad52::HIS VII-L::URA3-TG34-HOcs-LYS2 pif1-m2 rif1::KANMX* | This study |
| DDY3479 | S288C | *Mata-inc ura3-52 lys2-801 ade2-101 ochre trp1-Δ63 his3-Δ200 leu2-Δ1::GAL1:HO-LEU2 rad52::HIS VII-L::URA3-TG18-HOcs-LYS2 rif2::NATMX* | This study |
| DDY3480 | S288C | *Mata-inc ura3-52 lys2-801 ade2-101 ochre trp1-Δ63 his3-Δ200 leu2-Δ1::GAL1:HO-LEU2 rad52::HIS VII-L::URA3-TG18-HOcs-LYS2 pif1-m2 rif2::NATMX* | This study |
| DDY3481 | S288C | *Mata-inc ura3-52 lys2-801 ade2-101 ochre trp1-Δ63 his3-Δ200 leu2-Δ1::GAL1:HO-LEU2 rad52::HIS VII-L::URA3-TG34-HOcs-LYS2 rif2::NATMX* | This study |
| DDY3482 | S288C | *Mata-inc ura3-52 lys2-801 ade2-101 ochre trp1-Δ63 his3-Δ200 leu2-Δ1::GAL1:HO-LEU2 rad52::HIS VII-L::URA3-TG34-HOcs-LYS2 pif1-m2 rif2::NATMX* | This study |
| DDY3466 | S288C | *Mata-inc ura3-52 lys2-801 ade2-101 ochre trp1-Δ63 his3-Δ200 leu2-Δ1::GAL1:HO-LEU2 rad52::HIS VII-L::URA3-TG18-HOcs-LYS2 rif1::KANMX rif2::NATMX* | This study |
| DDY3467 | S288C | *Mata-inc ura3-52 lys2-801 ade2-101 ochre trp1-Δ63 his3-Δ200 leu2-Δ1::GAL1:HO-LEU2 rad52::HIS VII-L::URA3-TG18-HOcs-LYS2 pif1-m2 rif1::KANMX rif2::NATMX* | This study |
| DDY3468 | S288C | *Mata-inc ura3-52 lys2-801 ade2-101 ochre trp1-Δ63 his3-Δ200 leu2-Δ1::GAL1:HO-LEU2 rad52::HIS VII-L::URA3-TG34-HOcs-LYS2 rif1::KANMX rif2::NATMX* | This study |
| DDY3469 | S288C | *Mata-inc ura3-52 lys2-801 ade2-101 ochre trp1-Δ63 his3-Δ200 leu2-Δ1::GAL1:HO-LEU2 rad52::HIS VII-L::URA3-TG34-HOcs-LYS2 pif1-m2 rif1::KANMX rif2::NATMX* | This study |
| DDY3526 | S288C | *Mata-inc ura3-52 lys2-801 ade2-101 ochre trp1-Δ63 his3-Δ200 leu2-Δ1::GAL1:HO-LEU2 rad52::HIS VII-L::TG18-HOcs-LYS2 ura3::HPHMX cdc13::KANMX YEp-URA3-CDC13* | This study |
| DDY3527 | S288C | *Mata-inc ura3-52 lys2-801 ade2-101 ochre trp1-Δ63 his3-Δ200 leu2-Δ1::GAL1:HO-LEU2 rad52::HIS VII-L::URA3-TG18-HOcs-LYS2 pif1-m2 ura3::HPHMX cdc13::KANMX YEp-URA3-CDC13* | This study |
| DDY3528 | S288C | *Mata-inc ura3-52 lys2-801 ade2-101 ochre trp1-Δ63 his3-Δ200 leu2-Δ1::GAL1:HO-LEU2 rad52::HIS VII-L::TG18-HOcs-LYS2 ura3::HPHMX cdc13::KANMX YEp-URA3-CDC13* | This study |
| DDY3530 | S288C | *Mata-inc ura3-52 lys2-801 ade2-101 ochre trp1-Δ63 his3-Δ200 leu2-Δ1::GAL1:HO-LEU2 rad52::HIS VII-L::URA3-TG18-HOcs-LYS2 pif1-m2 ura3::HPHMX cdc13::KANMX YEp-URA3-CDC13* | This study |
| DDY3589 | S288C | *Mata-inc ura3-52 lys2-801 ade2-101 ochre trp1-Δ63 his3-Δ200 leu2-Δ1::GAL1:HO rad52::HIS VII-L::TG18-HOcs-LYS2 leu2::NAT ura3::HPHMX cdc13::KANMX YEp-URA3-CDC13 pRAD52-TRP* | This study |
| DDY3591 | S288C | *Mata-inc ura3-52 lys2-801 ade2-101 ochre trp1-Δ63 his3-Δ200 leu2-Δ1::GAL1:HO rad52::HIS VII-L::URA3-TG18-HOcs-LYS2 pif1-m2 leu2::NAT ura3::HPHMX cdc13::KANMX YEp-URA3-CDC13 pRAD52-TRP* | This study |
| DDY3534 | S288C | *Mata-inc ura3-52 lys2-801 ade2-101 ochre trp1-Δ63 his3-Δ200 leu2-Δ1::GAL1:HO-LEU2 rad52::HIS VII-L::TG18-HOcs-LYS2 ura3::HPHMX cdc13::KANMX pRS414-CDC13* | This study |
| DDY3536 | S288C | *Mata-inc ura3-52 lys2-801 ade2-101 ochre trp1-Δ63 his3-Δ200 leu2-Δ1::GAL1:HO-LEU2 rad52::HIS VII-L::TG18-HOcs-LYS2 ura3::HPHMX cdc13::KANMX pif1-m2 pRS414-CDC13* | This study |
| DDY3535 | S288C | *Mata-inc ura3-52 lys2-801 ade2-101 ochre trp1-Δ63 his3-Δ200 leu2-Δ1::GAL1:HO-LEU2 rad52::HIS VII-L::TG18-HOcs-LYS2 ura3::HPHMX cdc13::KANMX pRS414-cdc13-L91A* | This study |
| DDY3537 | S288C | *Mata-inc ura3-52 lys2-801 ade2-101 ochre trp1-Δ63 his3-Δ200 leu2-Δ1::GAL1:HO-LEU2 rad52::HIS VII-L::TG18-HOcs-LYS2 ura3::HPHMX cdc13::KANMX pif1-m2 pRS414-cdc13-L91A* | This study |
| DDY3538 | S288C | *Mata-inc ura3-52 lys2-801 ade2-101 ochre trp1-Δ63 his3-Δ200 leu2-Δ1::GAL1:HO-LEU2 rad52::HIS VII-L::TG34-HOcs-LYS2 ura3::HPHMX cdc13::KANMX pRS414-CDC13* | This study |
| DDY3540 | S288C | *Mata-inc ura3-52 lys2-801 ade2-101 ochre trp1-Δ63 his3-Δ200 leu2-Δ1::GAL1:HO-LEU2 rad52::HIS VII-L::TG34-* | This study |
|  |  | *HOcs-LYS2 ura3::HPHMX cdc13::KANMX pif1-m2*<br>*pRS414-CDC13* |  |
| DDY3539 | S288C | *Mata-inc ura3-52 lys2-801 ade2-101 ochre trp1-Δ63 his3-Δ200 leu2-Δ1::GAL1:HO-LEU2 rad52::HIS VII-L::TG34-HOcs-LYS2 ura3::HPHMX cdc13::KANMX pRS414-cdc13-L91A* | This study |
| DDY3541 | S288C | *Mata-inc ura3-52 lys2-801 ade2-101 ochre trp1-Δ63 his3-Δ200 leu2-Δ1::GAL1:HO-LEU2 rad52::HIS VII-L::TG34-HOcs-LYS2 ura3::HPHMX cdc13::KANMX pif1-m2*<br>*pRS414-cdc13-L91A* | This study |
| DDY3520 | S288C | *Mata-inc ura3-52 lys2-801 ade2-101 ochre trp1-Δ63 his3-Δ200 leu2-Δ1::GAL1:HO-LEU2 rad52::HIS VII-L::TG34-HOcs-LYS2 ura3::HPH cdc13::KANMX YEP24-CDC13*<br>*pRAD52* | This study |
| DDY3522 | S288C | *Mata-inc ura3-52 lys2-801 ade2-101 ochre trp1-Δ63 his3-Δ200 leu2-Δ1::GAL1:HO-LEU2 rad52::HIS VII-L::TG34-HOcs-LYS2 ura3::HPH cdc13::KANMX YEP24-CDC13*<br>*pRAD52 pif1-m2* | This study |
| DDY3528 | S288C | *Mata-inc ura3-52 lys2-801 ade2-101 ochre trp1-Δ63 his3-Δ200 leu2-Δ1::GAL1:HO-LEU2 rad52::HIS VII-L::TG34-HOcs-LYS2 ura3::HPH cdc13::KANMX YEP24-CDC13* | This study |
| DDY3530 | S288C | *Mata-inc ura3-52 lys2-801 ade2-101 ochre trp1-Δ63 his3-Δ200 leu2-Δ1::GAL1:HO-LEU2 rad52::HIS VII-L::TG34-HOcs-LYS2 ura3::HPH cdc13::KANMX YEP24-CDC13 pif1-m2* | This study |
| DDY3584 | S288C | *Mata-inc ura3-52 lys2-801 ade2-101 ochre trp1-Δ63 his3-Δ200 leu2-Δ1::GAL1:HO-LEU2 rad52::HIS VII-L::TG34-HOcs-LYS2 ura3::HPH leu2::NAT cdc13::KANMX YEP24-CDC13* | This study |
| DDY3586 | S288C | *Mata-inc ura3-52 lys2-801 ade2-101 ochre trp1-Δ63 his3-Δ200 leu2-Δ1::GAL1:HO-LEU2 rad52::HIS VII-L::TG34-HOcs-LYS2 ura3::HPH leu2::NAT cdc13::KANMX YEP24-CDC13 pif1-m2* | This study |
| DDY3589 | S288C | *Mata-inc ura3-52 lys2-801 ade2-101 ochre trp1-Δ63 his3-Δ200 leu2-Δ1::GAL1:HO-LEU2 rad52::HIS VII-L::TG34-HOcs-LYS2 ura3::HPH leu2::NAT cdc13::KANMX YEP24-CDC13 pRAD52* | This study |
| DDY3591 | S288C | *Mata-inc ura3-52 lys2-801 ade2-101 ochre trp1-Δ63 his3-Δ200 leu2-Δ1::GAL1:HO-LEU2 rad52::HIS VII-L::TG34-HOcs-LYS2 ura3::HPH leu2::NAT cdc13::KANMX YEP24-CDC13 pRAD52 pif1-m2* | This study |
| DDY3248 | S288C | *Mata-inc ura3-52 lys2-801 ade2-101 ochre trp1-Δ63 his3-Δ200 leu2-Δ1::GAL1:HO-LEU2 rad52::HIS VII-L::URA3-TG96-HOcs-LYS2* | This study |
| DDY3250 | S288C | *Mata-inc ura3-52 lys2-801 ade2-101 ochre trp1-Δ63 his3-Δ200 leu2-Δ1::GAL1:HO-LEU2 rad52::HIS VII-L::URA3-TG119-HOcs-LYS2* | This study |
| DDY3252 | S288C | *Mata-inc ura3-52 lys2-801 ade2-101 ochre trp1-Δ63 his3-Δ200 leu2-Δ1::GAL1:HO-LEU2 rad52::HIS VII-L::URA3-TG142-HOcs-LYS2* | This study |
| DDY3133 | S288C | *Mata-inc ura3-52 lys2-801 ade2-101 ochre trp1-Δ63 his3-Δ200 leu2-Δ1::GAL1:HO-LEU2 rad52::HIS VII-L::URA3-TG162-HOcs-LYS2* | This study |
| DDY3134 | S288C | *Mata-inc ura3-52 lys2-801 ade2-101 ochre trp1-Δ63 his3-Δ200 leu2-Δ1::GAL1:HO-LEU2 rad52::HIS VII-L::URA3-TG162-HOcs-LYS2 pif1-m2* | This study |
| DDY3529 | S288C | *Mata-inc ura3-52 lys2-801 ade2-101 ochre trp1-Δ63 his3-Δ200 leu2-Δ1::GAL1:HO-LEU2 rad52::HIS VII-L::TG34-HOcs-LYS2 ura3::HPH cdc13::KAN YEP24-CDC13* | This study |
| DDY3614 | S288C | DDY3529 *leu2::NATMX* | This study |
| DDY3531 | S288C | *Mata-inc ura3-52 lys2-801 ade2-101 ochre trp1-Δ63 his3-Δ200 leu2-Δ1::GAL1:HO-LEU2 rad52::HIS VII-L::TG34-HOcs-LYS2 ura3::HPH cdc13::KAN YEP24-CDC13 pif1-m2* | This study |
| DDY3615 | S288C | DDY3531 *leu2::NATMX* | This study |
| DDY3614 | S288C | *Mata-inc ura3-52 lys2-801 ade2-101 ochre trp1-Δ63 his3-Δ200 leu2-Δ1::GAL1:HO-LEU2 rad52::HIS VII-L::TG34-HOcs-LYS2 ura3::HPH cdc13::KAN leu2::NAT YEP24-CDC13* | This study |
| DDY3615 | S288C | *Mata-inc ura3-52 lys2-801 ade2-101 ochre trp1-Δ63 his3-Δ200 leu2-Δ1::GAL1:HO-LEU2 rad52::HIS VII-L::TG34-HOcs-LYS2 ura3::HPH cdc13::KAN leu2::NAT YEP24-CDC13 pif1-m2* | This study |
| DDY3768 | S288C | DDY3614 pRS425-*CDC13* | This study |
| DDY3769 | S288C | DDY3614 pRS425-*cdc13-L91A* | This study |
| DDY3770 | S288C | DDY3614 pRS425-*cdc13-F236S* | This study |
| DDY3771 | S288C | DDY3614 pRS425-*cdc13-Q256H* | This study |
| DDY3772 | S288C | DDY3614 pRS425-*cdc13-Q583K* | This study |
| DDY3773 | S288C | DDY3614 pRS425-*cdc13-I87N* | This study |
| DDY3774 | S288C | DDY3614 pRS425-*cdc13-Y758N* | This study |
| DDY3775 | S288C | DDY3614 pRS425-*cdc13-H12R* | This study |
| DDY3776 | S288C | DDY3614 pRS425-*cdc13-E556V/N567D* | This study |
| DDY3777 | S288C | DDY3614 pRS425-*cdc13-F728I* | This study |
| DDY3778 | S288C | DDY3614 pRS425-*cdc13-E252K* | This study |
| DDY3779 | S288C | DDY3614 pRS425-*cdc13-P235S* | This study |
| DDY3780 | S288C | DDY3614 pRS425-*cdc13-S255A* | This study |
| DDY3781 | S288C | DDY3614 pRS425-*cdc13-K50Q* | This study |
| DDY3782 | S288C | DDY3614 pRS425-*cdc13-F237V* | This study |
| DDY3783 | S288C | DDY3615 pRS425-*CDC13* | This study |
| DDY3784 | S288C | DDY3615 pRS425-*cdc13-L91A* | This study |
| DDY3785 | S288C | DDY3615 pRS425-*cdc13-F236S* | This study |
| DDY3786 | S288C | DDY3615 pRS425-*cdc13-Q256H* | This study |
| DDY3787 | S288C | DDY3615 pRS425-*cdc13-Q583K* | This study |
| DDY3788 | S288C | DDY3615 pRS425-*cdc13-I87N* | This study |
| DDY3789 | S288C | DDY3615 pRS425-*cdc13-Y758N* | This study |
| DDY3790 | S288C | DDY3615 pRS425-*cdc13-H12R* | This study |
| DDY3791 | S288C | DDY3615 pRS425-*cdc13-E556V/N567D* | This study |
| DDY3792 | S288C | DDY3615 pRS425-*cdc13-F728I* | This study |
| DDY3793 | S288C | DDY3615 pRS425-*cdc13-E252K* | This study |
| DDY3794 | S288C | DDY3615 pRS425-*cdc13-P235S* | This study |
| DDY3795 | S288C | DDY3615 pRS425-*cdc13-S255A* | This study |
| DDY3796 | S288C | DDY3615 pRS425-*cdc13-K50Q* | This study |
| DDY3797 | S288C | DDY3615 pRS425-*cdc13-F237V* | This study |

